# Error minimization and specificity could emerge in a genetic code as by-products of prebiotic evolution

**DOI:** 10.1101/2021.05.14.444235

**Authors:** Evan Janzen, Yuning Shen, Ziwei Liu, Celia Blanco, Irene A. Chen

## Abstract

The emergence of the genetic code was a major transition in the evolution from a prebiotic RNA world to the earliest modern cells^1^. A prominent feature of the standard genetic code is error minimization, or the tendency of mutations to be unusually conservative in preserving biophysical features of the amino acid^2–6^. While error minimization is often assumed to result from natural selection, it has also been speculated that error minimization may be a by-product of emergence of the genetic code^3^. During establishment of the genetic code in an RNA world, self-aminoacylating ribozymes would enforce the mapping of amino acids to anticodons. Here we show that expansion of the genetic code, through co-option of ribozymes for new substrates, could result in error minimization as an emergent property. Using self-aminoacylating ribozymes previously identified during an exhaustive search of sequence space^7^, we measured the activity of thousands of candidate ribozymes on alternative substrates (activated analogs for tryptophan, phenylalanine, leucine, isoleucine, valine, and methionine). Related ribozymes exhibited preferences for biophysically similar substrates, indicating that co-option of existing ribozymes to adopt additional amino acids into the genetic code would itself lead to error minimization. Furthermore, ribozyme activity was positively correlated with specificity, indicating that selection for increased activity would also lead to increased specificity. These results demonstrate that by-products of the evolution and functional expansion of a ribozyme system could lead to adaptive properties of a genetic code. Such ‘spandrels’ could thus underlie significant prebiotic developments.

## Introduction

The origin of life is believed to have progressed through an RNA World in which ribozymes catalyzed critical biochemical reactions^8,9^. In principle, ribozymes performing new functions could arise either by chance, or by adaptation of pre-existing ribozymes having promiscuous activities. Co-option of a pre-existing sequence (i.e., exaptation) is a well-established mechanism for evolutionary innovation^10–15^. Gene duplication coupled with co-option could lead to a more complex system as the ribozymes adopt additional substrates^16^. However, the degree to which the evolution of complex systems in the RNA World would rely on chance vs. co-option is unclear^17^.

The genetic code of protein translation is one of the most complex products of the RNA World, and its emergence is considered a ‘major evolutionary transition’^1^. In modern biology, the mapping of specific codons to their cognate amino acids is assured through the aminoacylation of tRNAs by aminoacyl-tRNA synthetase (aaRS) proteins^18–20^. However, during the emergence of protein translation itself, these functions were presumably performed by ribozymes. Indeed, evolutionary analysis of the aaRS proteins indicates that these enzymes evolved after the establishment of a primitive genetic code^21–25^ and have heterogeneous genetic origins^26^. Several ribozymes catalyzing aminoacylation reactions have been discovered by *in vitro* selection, including self-aminoacylating RNAs^7,27–31^. Such ribozymes could serve as precursors to the aaRS/tRNA encoding system.

A well-documented feature of the standard genetic code is robustness to errors, i.e., that non-synonymous point mutations tend to result in amino acid substitutions that conserve biophysical properties^2–6^. This ‘error minimization’ confers a clear selective advantage as it reduces the deleterious impact of mutations on the resultant protein^32,33^. However, the standard genetic code does not appear to be particularly optimal with respect to error minimization^34–37^. This raises a fundamental open question about the origin of error minimization, namely, whether error minimization of the standard genetic code is the product of natural selection, or a serendipitous by-product of the evolution of protein translation^3^. In other words, in contrast to direct natural selection for error minimization, it is possible that expansion of an early version of the code, initially comprising a small number of amino acids, to the full set of 20 amino acids, involved an evolutionary mechanism that happened to conserve the biophysical character of the amino acids^38,39^.

In this work, we evaluate the evolutionary potential of self-aminoacylating ribozymes to adopt new amino acid substrates. We previously used *in vitro* selection and high-throughput sequencing to exhaustively search RNA sequence space (21 nt) for self-aminoacylating ribozymes^7^. These ribozymes were originally selected to react with biotinyl-Tyr(Me)-oxazolone (BYO), a chemically activated amino acid. The 5(4*H*)-oxazolones and related *N*-carboxyanhydrides can be made abiotically under prebiotically plausible conditions ^40–48^. Three distinct, evolutionarily unrelated catalytic motifs had been discovered from the exhaustive search. Here we determine the co-option potential of these ribozymes, by measuring the activity of all single- and double- mutants of five ribozymes, representing the three catalytic motifs, for six alternative substrates, using a massively parallel assay (*k*-Seq^7,49^). This assay and related techniques leverage high-throughput sequencing to measure the activity of thousands of candidate sequences in a mixed pool^50–53^. The six substrates (analogs of tryptophan, phenylalanine, leucine, isoleucine, valine, and methionine) represent a range of sizes and biophysical classes (aromatic, aliphatic, sulfur-containing), as well as supposed early (Leu, Ile, Val) and late (Trp, Phe, Met) incorporations into the genetic code^54–58^. Our findings indicate extensive opportunities for co-option to incorporate new substrates into the system. In addition, we describe two major by-products of evolution of these ribozymes. First, a positive correlation between activity and specificity was observed, indicating that greater specificity would be a by-product of selection for greater activity. Second, related ribozymes react with biophysically similar amino acids, suggesting that expansion of the code by co-option would incorporate a biophysically similar amino acid into the system, with error minimization arising as a by-product. Such effects could favor the emergence of a complex biochemical system.

## Results

### Aminoacylation substrates and design of the ribozyme pool

To investigate whether ribozymes previously selected for aminoacylation with BYO (tyrosine analog) would react with substrates having other aminoacyl side chains, six additional biotinyl-aminoacyl oxazolones were synthesized for analysis (Figure 1A): tryptophanyl (BWO), phenylalanyl (BFO), leucyl (BLO), isoleucyl (BIO), valyl (BVO), and methionyl (BMO). Compounds were synthesized using previously described methods^7^ and verified by NMR spectroscopy (see Methods). An initial test by a gel shift assay at high substrate concentration (500 μM) indicated that each oxazolone served as substrate for at least one ribozyme tested, although the two tested ribozymes (S-1A.1-a and S-2.1-a) differed in selectivity (Figure 1B). To study the cross-reactivity of these ribozymes and their mutants systematically, pools of sequence variants were designed to explore the sequence space around the major ribozyme families obtained from the selection on BYO (Table S1). The ribozyme families chosen for testing include all of the previously discovered motifs (Motifs 1, 2, and 3), specifically the two most abundant families containing Motif 1 (Family 1A.1 and 1B.1) and Motif 2 (Family 2.1 and 2.2), as well as the only family identified from Motif 3 (Family 3.1). These ribozyme families had been discovered during an exhaustive search of sequence space varying a central 21-mer region, and sequences from these motifs had comprised ~80% of the selected pool^7^. Sequencing of the variant pool showed that it included 13.5% of the unique sequences from the originally selected pool (having abundance ≥10^−6^). Thus, the variant pool, based on these five ribozyme families, was designed to be representative of ribozymes having aminoacylation activity.

**Figure 1.**
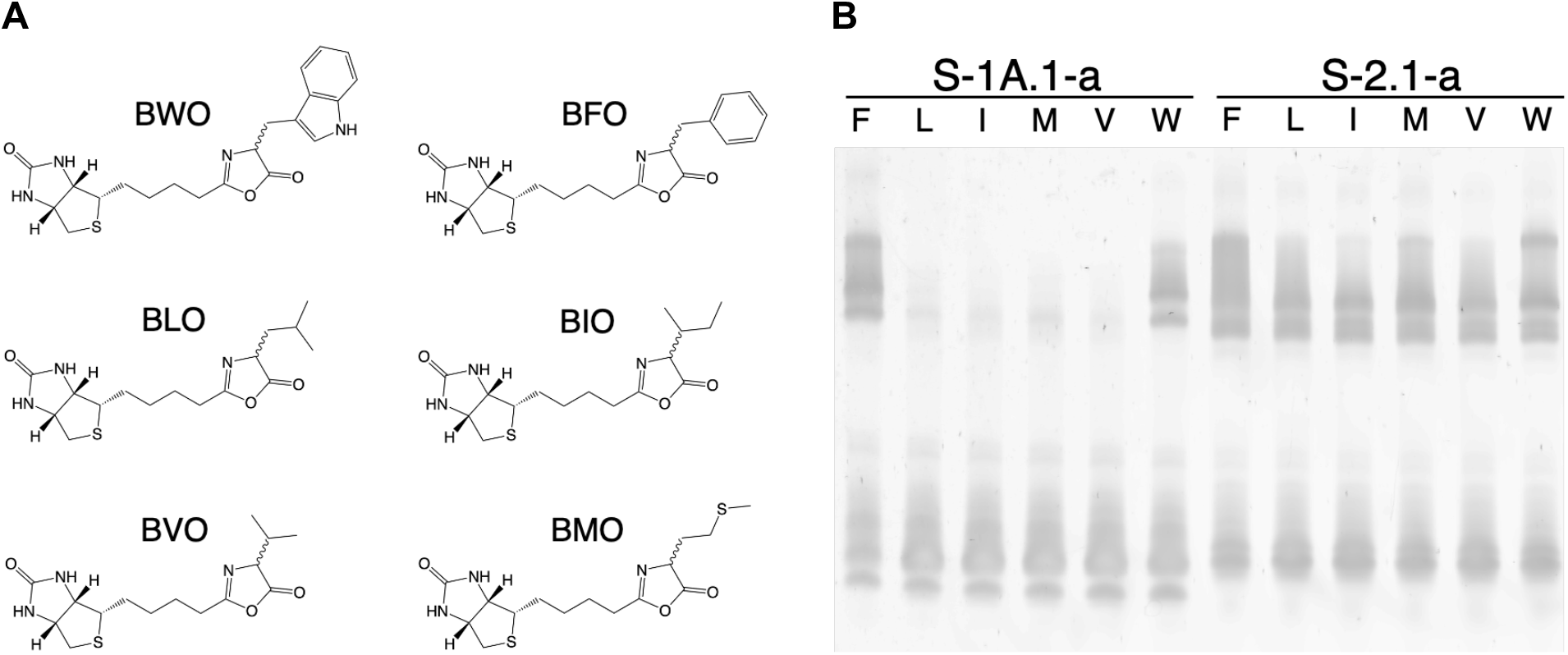
Aminoacylation activity of two ribozymes with BXO substrates. A) Biotinyl aminoacyl oxazolones (BXO) used in this study: tryptophanyl (BWO), phenylalanyl (BFO), leucyl (BLO), isoleucyl (BIO), valyl (BVO), and methionyl (BMO). B) Aminoacylation activity of two ribozymes (S-1A.1-a, the center of Family 1A.1, and S-2.1-a, the center of Family 2.1) with BXO substrates analyzed by streptavidin gel shift (X = F, L, I, M, V, or W, as indicated). Reactions were conducted for 90 min at 500 μM BXO. The reacted RNA is detected by its slower migration through the gel due to complexation with streptavidin. Multiple bands may be caused by the presence of multiple conformers or streptavidin oligomers.

Because the ribozymes had been identified through selection with substrate BYO, it was possible that entirely new ribozyme families might react with different BXO substrates. To assess this possibility, *in vitro* selections for self-aminoacylating ribozymes were performed for two of the new substrates (BFO and BLO), starting from libraries with completely random 21-mer variable regions. These selections followed a process identical to the original selection with the exception of the substrate compound. All families found in the BFO and BLO selections had already been identified in the earlier BYO selection (Figure S1). Interestingly, selection with BLO resulted predominantly in sequences containing Motif 2, consistent with the low activity of a Family 1A.1 ribozyme on BLO observed in the gel shift assay (Figure 1B). These results indicate that the designed pool of variants would probe the major motifs of the active sequence space for these substrates.

### Cross-reaction of self-aminoacylating ribozymes with alternative side chains

Sequences in the ribozyme variant pool were assayed for activity on each alternative substrate in a massively parallel format by kinetic sequencing (*k*-Seq)^7,49,59^. During *k*-Seq, a pool containing thousands of candidate ribozymes is reacted with a substrate at multiple concentrations. The reacted molecules, having been biotinylated through reaction, are isolated by streptavidin binding and then sequenced on the Illumina platform. Quantitation of the reacted fraction allows fitting to a kinetic model to determine ribozyme activity. Data obtained from this method correlate well with traditional biochemical assays, provided a sufficient number of sequencing counts, and confidence intervals of the measurements are obtained by experimental replicates and bootstrapping^49^. In each *k*-Seq experiment here, one of six BXO (X = W, F, L, I, V, or M) substrates was tested to measure reaction kinetics for sequences in the pool. Samples were exposed to substrate concentrations from 2 to 1250 μM in triplicate. Reaction data were fit to a pseudo-first-order kinetic model 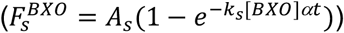, with maximum reaction amplitude *A_s_* and rate constant *k_s_* for sequence *s*, where 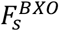 is the fraction of RNA that is aminoacylated with substrate BXO, [BXO] is the initial substrate concentration, *t* is the reaction time (90 min), and *α* is the coefficient accounting for substrate hydrolysis during the reaction. Although data over a fixed concentration range are inadequate for separately estimating *k_s_* and *A_s_* for low activity ribozymes, the product *k_s_A_s_* can be accurately estimated across a wide range of activities, due to the inverse correlation of *k_s_* and *A_s_* during curve fitting^7,49^ (Figure S2). The product *k_s_A_s_* reflects ribozyme activity at non-saturating conditions and was used in the following analyses. The data yielded *k_s_A_s_* estimates for a total of 9,770 sequences, encompassing five family wild-type sequences and a complete set of both single and double mutants related to the five wild-type ribozymes (Figure S3).

*k*-Seq measures the combination of catalyzed and non-catalyzed (background) reactions. To determine catalytic enhancement, i.e., the ratio of catalyzed to background reaction rates, we measured the rate of the background reaction for BFO by gel shift assay with the randomized RNA library. The background rate was 0.55 ± 0.18 M^−1^min^−1^ (μ ± σ), which is similar to that measured previously for BYO (0.65 ± 0.28 M^−1^min^−1^)^7^. Comparing to the frequency distribution of *k_s_A_s_* measured by *k*-Seq (Figure S4, Table S2), the measured background rate was found to correspond to the center of a low-activity peak, indicating that this peak represented a background of catalytically inactive, or nearly inactive, mutants. This is consistent with observations that individual Motif 1 ribozymes display little activity with some substrates at high concentration when analyzed by a gel-shift assay (Figure 1B). The low-activity peak was therefore used as an internal control in *k*-Seq, and the effective background reaction rate (*k*_0_*A*_0_) of each substrate was estimated as the center of this peak. *k_s_A_s_* values for sequences reacted with each substrate were normalized by the corresponding *k*_0_*A*_0_ to obtain the catalytic enhancement above background, or *r_s_* (defined as *r_s_* = *k_s_A_s_*/*k*_0_*A*_0_ for each sequence *s*).

The *r_s_* values obtained from the *k*-Seq experiments revealed that all tested families contained sequences which displayed some activity on a new substrate or on multiple new substrates (Figure 2). Details of the frequency distribution of catalytic enhancement depended on both the aminoacyl side chain of the substrate as well as the ribozyme family. The distribution of sequences in Families 1A.1, 1B.1, and 3.1 could be characterized as containing a peak centered around background activity accompanied by a long, high-activity tail, particularly with BWO and BFO (Figure S5). In contrast, the distributions of Families 2.1 and 2.2 displayed distinct peaks at higher activity, with bimodality apparent in some cases (especially for Family 2.1). This indicated a higher tolerance for mutations in Families 2.1 and 2.2 than in 1A.1, 1B.1, and 3.1, as mutant sequences were less likely to exhibit substantial detrimental effects.

**Figure 2.**
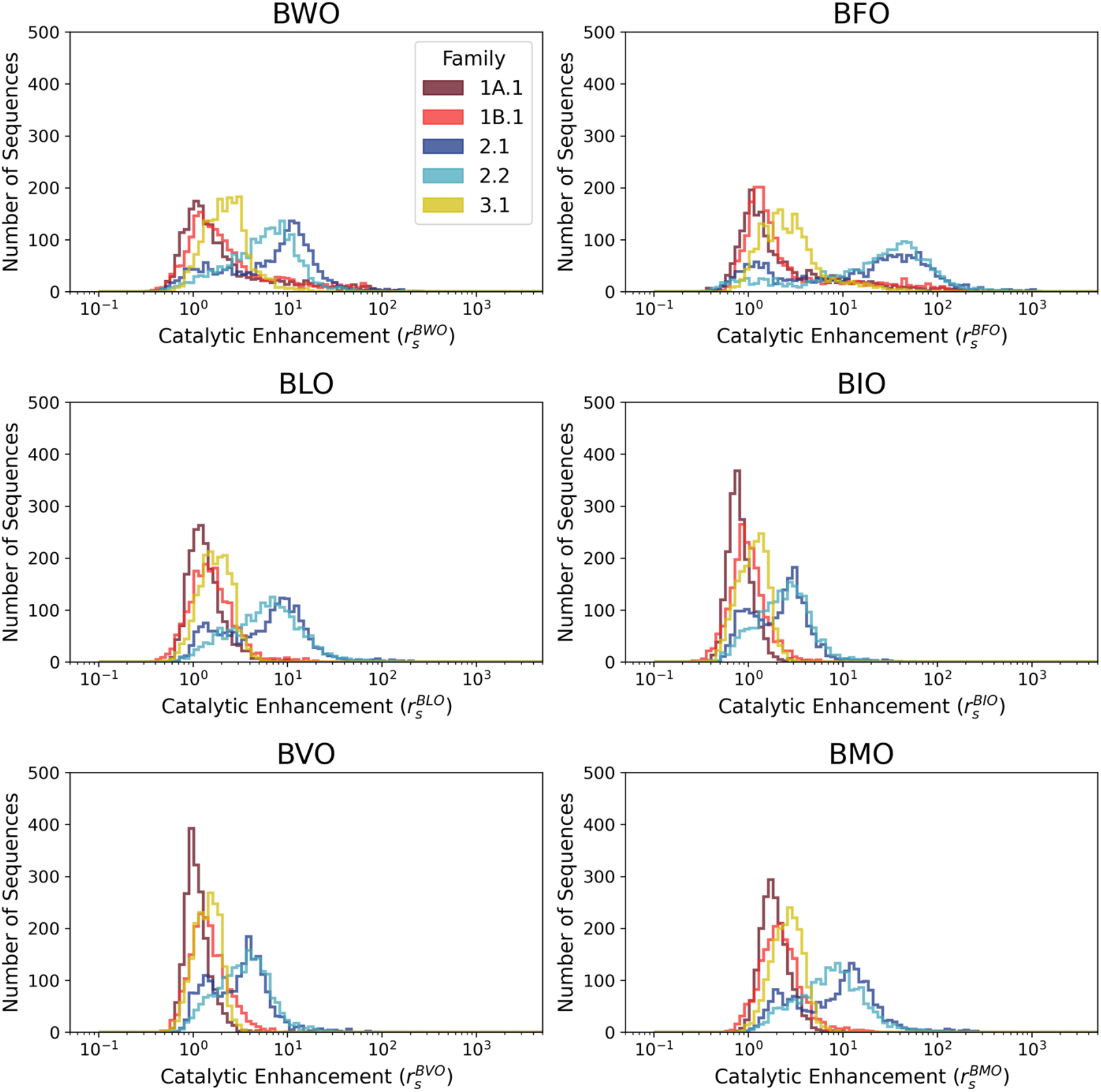
Catalytic enhancement of ribozyme families for different substrates. Histograms of catalytic enhancement values (*r_s_* = *k_s_A_s_/k_0_A_0_*) with each BXO substrate, measured by *k*-Seq, for ribozymes in Family 1A.1, 1B.1, 2.1, 2.2, and 3.1. While many ribozyme mutants in Motif 2 families have activity on each substrate tested, many ribozyme sequences containing Motif 1 or 3 are inactive.

### Ribozyme families distinguish different biophysical features of substrate side chains

To assess the activity and specificity of individual ribozymes for each substrate, catalytic enhancement values for different substrates were compared in a pairwise fashion (Figures 3 and S6). All families displayed a high degree of correlation among activities for non-aromatic amino acid analogs (BLO (Leu), BIO (Ile), BVO (Val), and BMO (Met)) and also between activities for the two aromatic analogs (BWO (Trp) and BFO (Phe)) (Figure 4A). The high correlations indicated that few sequences exhibit large activity differences between amino acids within the same biophysical class.

**Figure 3.**
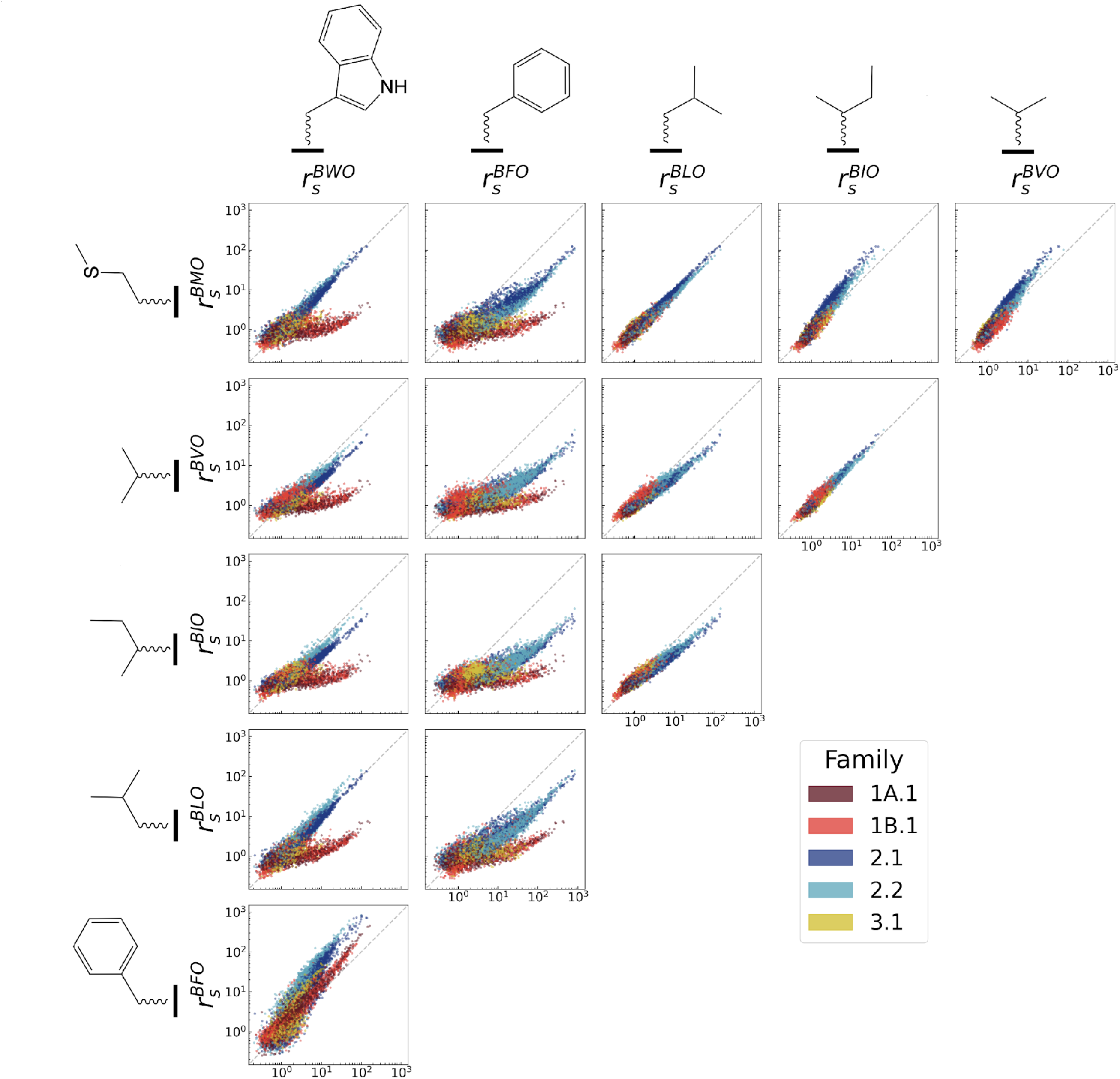
Pairwise comparisons of ribozyme activity on different substrates. A) Pairwise comparisons of catalytic enhancement (*r_s_*) for individual ribozyme sequences with each BXO substrate. Dashed gray line indicates the identity line. Substrates are ordered by hydrophilicity^60^. See Figure S6 for error bars and mutant order for each family.

**Figure 4.**
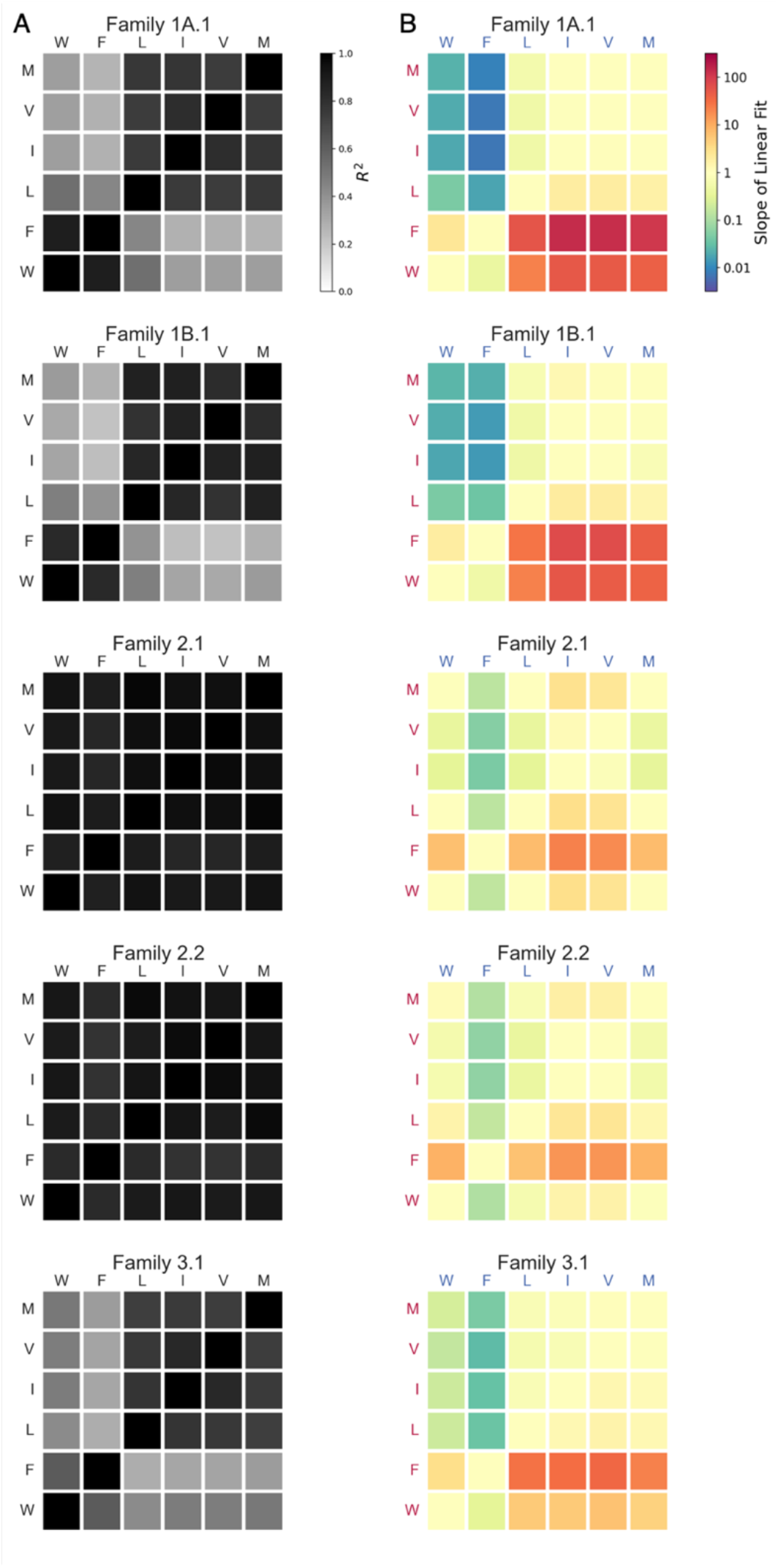
Ribozyme substrate preferences and correlations of activity. A) Heat maps of coefficient of determination (R^2^) for pairwise comparisons in Figure 3. B) Heat maps for slopes of linear regression fits for pairwise comparisons in Figure 3. Slope > 1 indicates a preference for the substrate on the y-axis; slope < 1 indicates a preference for the substrate on the x-axis.

However, when comparing amino acids of different classes (i.e., aromatic vs. non-aromatic), strong correlations were only observed for Families 2.1 and 2.2, indicating that the effects of mutations in Motif 2 ribozymes tend to be relatively independent of the side chain. In contrast, Families 1A.1, 1B.1, and 3.1 showed substantially lower activity with non-aromatic side chains (Figure 3), resulting in lower correlations between activity on aromatic and non-aromatic side chains (Figure 4A). These preferences were also captured by the slopes on the correlation plots (Figure 4B), which confirm that Motif 1 ribozymes strongly favor aromatic side chains, while Motif 2 ribozymes demonstrate less pronounced preferences, and Motif 3 ribozymes display an intermediate strength of preference. While less pronounced than for Motif 1, some preferences were still observed for Motif 2 ribozymes, in which BFO was most preferred, BMO, BWO and BLO were weakly preferred, and BVO and BIO were disfavored. Interestingly, BVO and BIO, in contrast to the other side chains, are both branched at the β carbon position. For Family 3.1, BFO was preferred over BWO, and all non-aromatic substrates were similarly disfavored. The differences observed between trends characterizing the separate ribozyme motifs suggest differences in the recognition mechanisms among Motifs 1, 2, and 3. Nevertheless, all ribozyme families display some preferences that correspond to biophysical features of the side chains.

### Substrate specificity is positively correlated with activity

To probe the relationship between catalytic activity and substrate specificity, we used two measures of specificity. First, as a general measure of substrate specificity for each sequence, we adapted the ‘promiscuity index’^61^. This metric 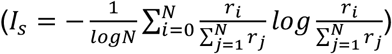 is a normalized entropy which describes the evenness of rates across different substrates. The promiscuity index *I_s_* ranges from 0 to 1, such that sequences that are completely promiscuous have *I_s_* = 1 and sequences completely specific to one substrate have *I*_s_ = 0. Promiscuity was observed to decrease as overall activity increased for all families (Figures 5 and S7).

**Figure 5.**
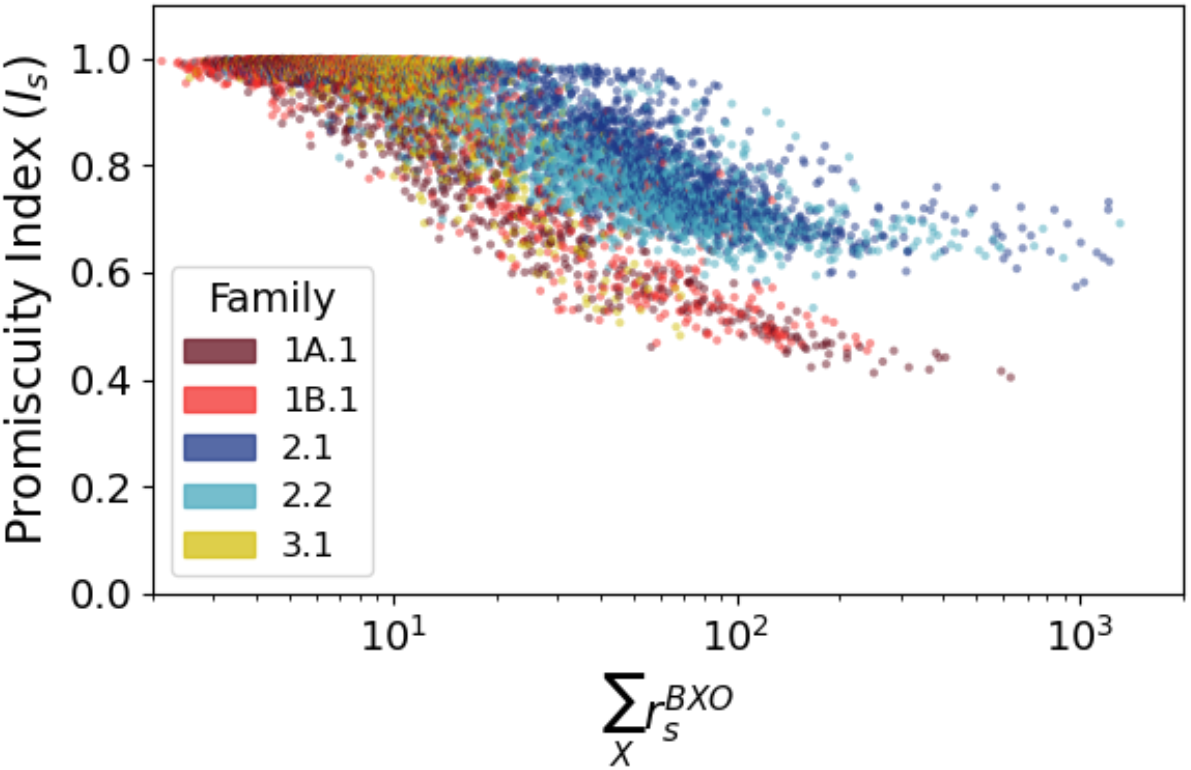
Relationship between activity and promiscuity. Promiscuity index values for each sequence as a function of total activity (sum of activities with all tested substrates). The general trend indicates that specificity increases (promiscuity decreases) as overall activity increases.

Second, since ribozymes in some families displayed preferential activity with aromatic amino acids compared to non-aromatic amino acids, we calculated the relative preference for aromatic substrates as 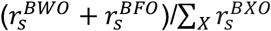. This ‘aromatic preference’ ratio reflects the proportion of ribozyme products that would have aromatic side chains in a reaction containing all six substrates at equal, sub-saturating concentration (Figure S8). Both the aromatic preference and the promiscuity index showed that the total activity of a ribozyme was positively correlated with specificity (positively correlated with aromatic preference and negatively correlated with promiscuity index; Table 1).

**Table 1.** Correlations between overall catalytic activity and specificity for each ribozyme family (Pearson’s *r* and Spearman’s ρ; *n* = 1954, *p*-values < 10^−95^ in each case).

| Family | Promiscuity Index |  | Aromatic Preference |  |
| --- | --- | --- | --- | --- |
| | $r$ | $\rho$ | $r$ | $\rho$ |
| **1A.1** | -0.696 | -0.647 | 0.554 | 0.711 |
| **1B.1** | -0.839 | -0.502 | 0.738 | 0.477 |
| **2.1** | -0.535 | -0.888 | 0.452 | 0.911 |
| **2.2** | -0.538 | -0.866 | 0.445 | 0.865 |
| **3.1** | -0.814 | -0.462 | 0.749 | 0.513 |

### Abundance of opportunities for co-option for alternative substrates

To quantify the frequency of sequences with potential for co-option, we categorized sequences as active or inactive using a catalytic enhancement threshold *r_t_*. Sequences below this threshold are considered to be nearly inactive, being close to the background rate (see above). An activity threshold of *r_t_* = 5 was chosen for two reasons. First, this threshold is two-fold more than the estimated 95% range for background activity (Figure S4, Table S2), so values of *r_s_* > 5 are statistically significantly greater than the normalized background rate. Second, increasing the rate of reaction by a factor of 5 is potentially significant in a prebiotic context, as abundances are expected to depend exponentially on relative fitness. Using this threshold, ribozymes that were active on more than one substrate were considered capable of exaptation.

Consistent with the observation that sequences in Families 2.1 and 2.2 displayed a high level of correlation of activities among all tested substrates, these families also yielded abundant opportunities for co-option, with most sequences being active with at least two substrates (1029 sequences in Family 2.1; 853 sequences in Family 2.2), and many active with all six tested substrates (Figure 6). In contrast, Families 1A.1, 1B.1, and 3.1, which contain more inactive sequences and generally preferred aromatic amino acids, yielded fewer exaptation opportunities, with most sequences accepting only one (or zero) substrates. Of sequences capable of exaptation in Families 1A.1, 1B.1, and 3.1, most were only active with two substrates. Nevertheless, even in these families, >2% of sequences accepted 2 or more substrates (254 sequences in Family 1A.1, 278 sequences in Family 1B.1, and 43 sequences in Family 3.1).

**Figure 6.**
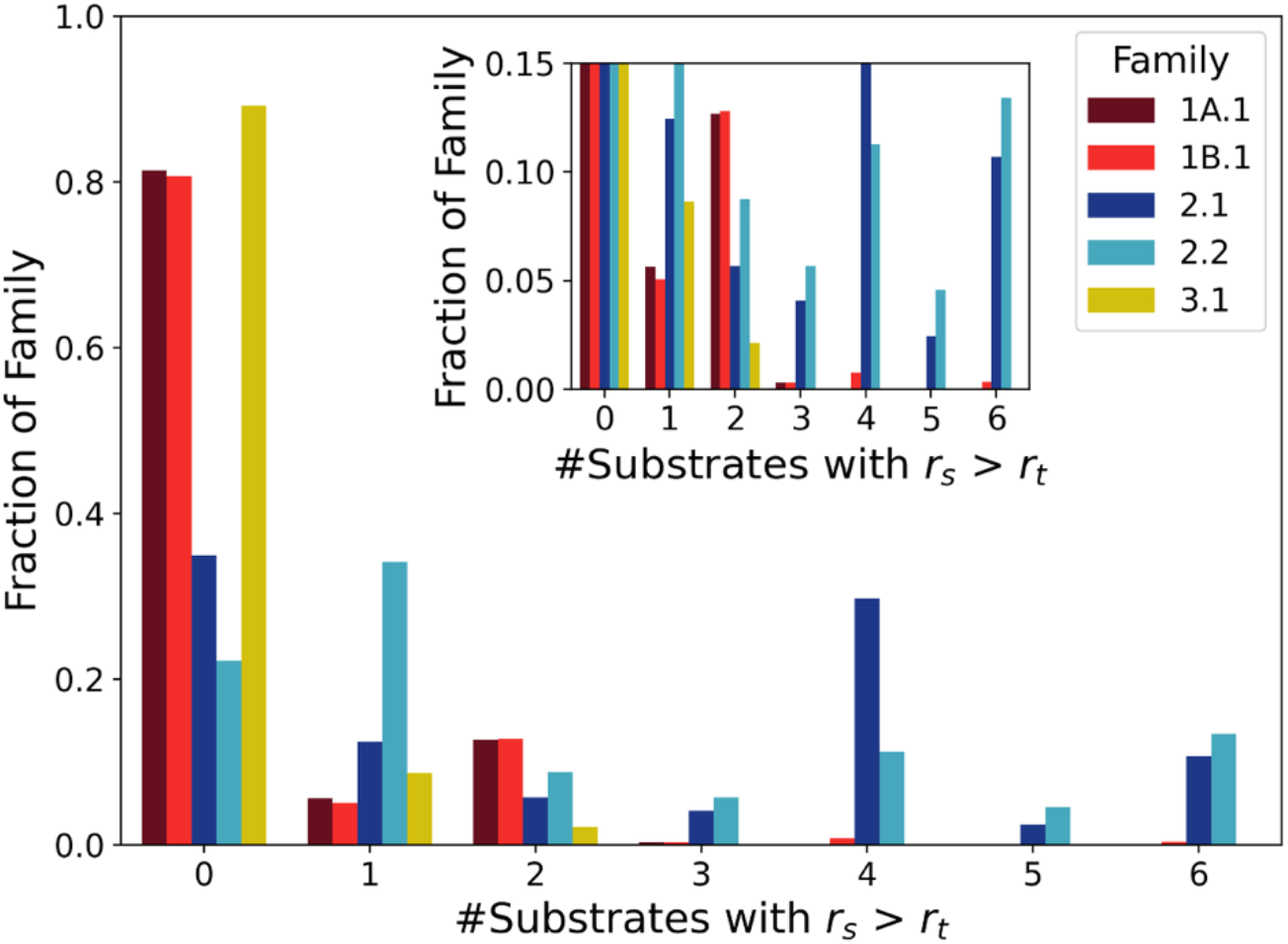
Ribozyme sequences with co-option potential. The frequency distribution of the fraction of unique sequences in each family (y-axis) that is active on a given number of substrates (x-axis). Activity on 2 or more substrates indicates potential for co-option. While Motif 2 sequences (Families 2.1 and 2.2) show a higher abundance of sequences active on more substrates, all families possess some co-option potential. Inset shows an enlargement of the low *y*-value region of the plot.

### Optimization of co-opted function on the fitness landscape

The sequences identified as presenting opportunities for co-option are active on two (or more) substrates, but may not be optimally active on either. To determine how readily co-option might lead to an optimally active sequence on a given substrate through evolution over the fitness landscape, we investigated the connectivity of optimal sequences (i.e., fitness peaks) for each substrate within the fitness landscape defined by each substrate, for each ribozyme family. With the exception of Family 3.1, the substrate peaks (highest *r_s_*) for each family were accessible to one another by evolutionary pathways proceeding through single mutations, while maintaining some activity (i.e., maintaining 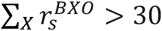, in analogy to *r_t_* = 5 for 6 substrates) (Figure 7). Family 3.1 was unique among families, in that the few co-optable sequences active on non-aromatic substrates were isolated in sequence space from the larger number of aromatic-preferring ribozymes.

**Figure 7.**
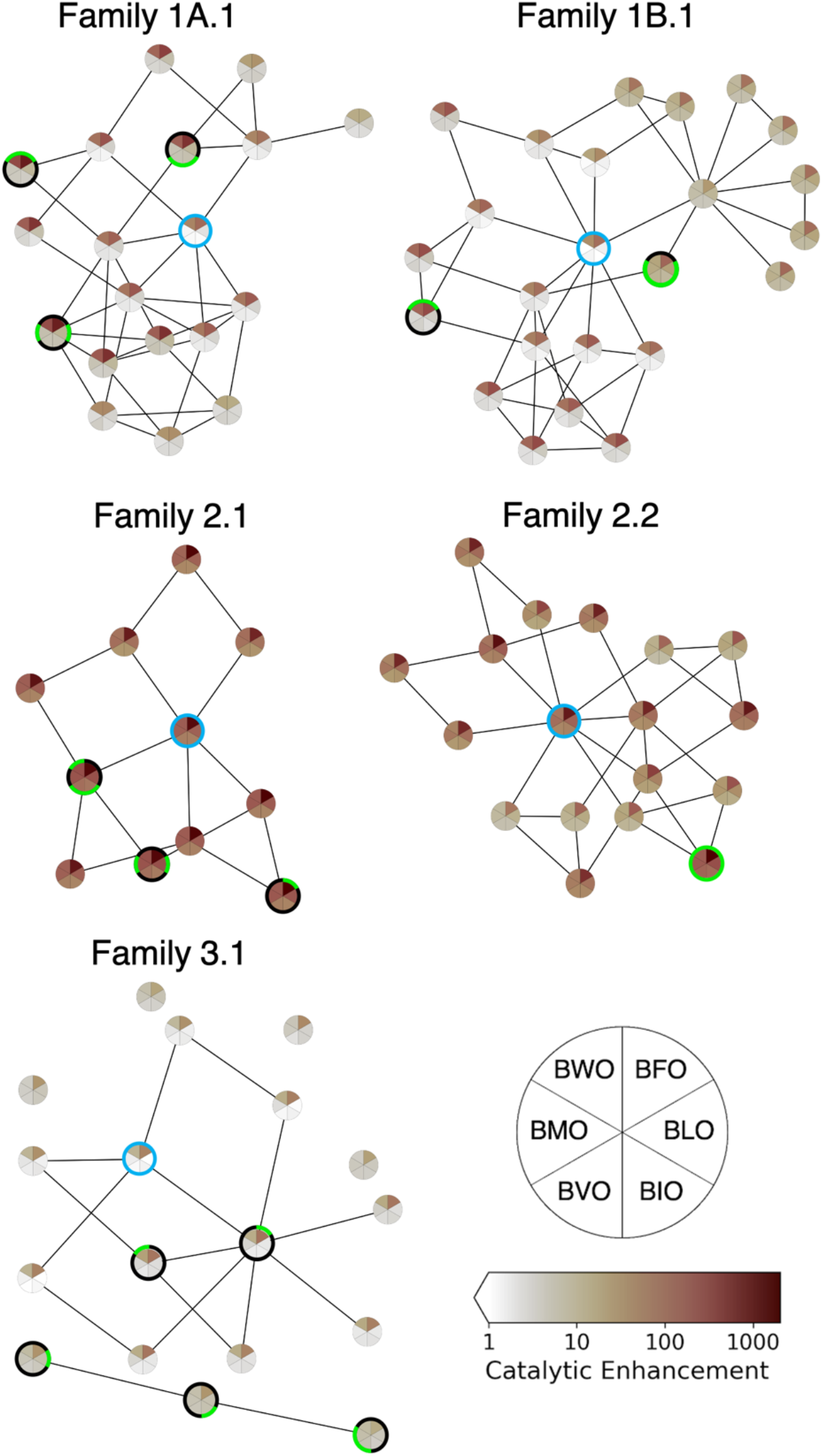
Evolutionary pathways for optimization from potential co-option points on the fitness landscape. Each circular ‘pie’ represents a single sequence, whose catalytic enhancement for each substrate is shown by sector shading according to the heat map legend. For each family, the wild-type and the ribozymes having the six highest catalytic enhancements for each substrate are included. The wild-type sequence in each family is highlighted by a blue circle; the most active sequence for each substrate is indicated by a green sector outline for the substrate. Among the set of high-activity sequences, every pair of sequences for which Hamming distance *d* = 2 was examined to identify intervening sequences (*d* = 1 to both sequences of the pair) having substantial overall activity 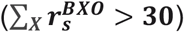. The intervening sequences are also shown in the plot. Lines connect sequences where *d* = 1. Sequences and catalytic enhancement values are given in Table S3.

## Discussion

The genetic code is an ideal platform for studying co-option in ribozyme evolution, as aminoacylations by the 20 biogenic amino acids represent naturally distinct functions. The genetic code itself is thought to have been established during the RNA World, in which ribozymes catalyzed aminoacylation^21–24^. Here we determined the activities of self-aminoacylating ribozyme families with several activated amino acid substrates. These ribozymes were originally discovered by exhaustive *in vitro* selection over sequence space (21 nt random region flanked by constant regions)^7^, and thus their properties are expected to be a reasonable model for self-aminoacylating ribozymes. Each tested family contained dozens or hundreds of ribozyme sequences that could utilize multiple substrates, often with high correlations in activity between substrates. In addition, the optimally active sequences with each substrate were closely connected in sequence space in four of the five families, demonstrating high evolvability and optimization potential between functions. This highlights the potential for ribozymes with activity for a selected substrate to adopt other amino acid substrates. In an RNA World scenario, this process could be beneficial for expanding metabolic chemical space and incorporating new compounds into increasingly complex systems.

While all families displayed substantial potential for adopting new substrates through co-option, ribozyme families differed in substrate preference and overall activity. Namely, Families 1A.1, 1B.1, and 3.1 contained relatively few active ribozymes, and these tended to display strong preference for aromatic amino acid side chains, although some sequences in these families were more promiscuous. The families in Motif 1 followed the general preference order of F,W > M,L,I,V, and the Motif 3 family followed the general preference order of F > W > M,L,I,V. Thus, these ribozymes appear to distinguish aromatic and non-aromatic side chains. On the other hand, Families 2.1 and 2.2 contained many sequences with high activity on all tested substrates, and also tended to prefer BFO. The families in Motif 2 followed the general preference order of F > M,W,L > I,V. This preference order suggests that Motif 2 ribozymes prefer the aromatic side chains, and are also subject to steric constraints, as they prefer F over W and also prefer L (non-branched β-carbon) over I and V (branched β-carbon). Given that these ribozymes were not selected for specificity (i.e., no counter-selections or negative selections), these preferences reflect inherent biophysical and structural features of the RNA interactions with different side chains.

The evolution of error minimization in the standard genetic code has been a subject of extensive theoretical and analytical study stemming from the realization that the code is unusually conservative in light of mutations. Since error minimization has adaptive value, a prevalent and intuitive view is that this property arose through natural selection^5,6,62^. However, an alternative view is that this trait emerged as a by-product during the initial expansion of the genetic code^36,37,39^. For example, it has been suggested that duplication of aminoacyl-tRNA synthetases would lead to emergence of a conservative pairing, as the tRNA and amino acid would be similar to the ancestral versions^63^. Since the catalytic elements of the earliest protein translation machinery were presumably composed of RNA, and indeed, phylogenetic evidence suggests that the genetic code predates aminoacyl-tRNA synthetases, a similar logic suggests that code expansion in the RNA World would have a tendency to conserve biophysical features of the substrate^38,39^. Using our experimental system of self-aminoacylating RNAs, we found that all ribozymes showed preferences for certain biophysical features, being particularly sensitive to aromaticity and branching in the side chain. Thus, co-option of these ribozymes would produce an association between these biophysical features and the RNA sequence, possibly including the primitive anticodon region. While the self-aminoacylating ribozymes studied here are a model system and not expected to recapitulate the evolution of the existing standard genetic code, these results illustrate the feasibility of the general principle that ribozyme co-option to incorporate new amino acid substrates would lead to error minimization as a by-product of expansion of the genetic code.

Substrate preferences were amplified with increasing activity, resulting in a positive correlation between activity and substrate specificity. Previous research on the relationship between activity and specificity has noted intuitively appealing trade-offs between these two properties in some systems^64–70^, as may be caused by ground-state discrimination in enzymes. In contrast, the results seen here indicate a positive correlation between catalytic activity and substrate specificity, instead reminiscent of enzymes that employ transition-state discrimination^69,71^. The evolutionary consequence of the positive activity-specificity correlation is that natural selection for greater activity would also lead to greater substrate specificity, as a by-product. At the same time, given the prevalence of promiscuous sequences and the short evolutionary pathways among optimal sequences for different substrates, new substrate specificities would still be accessible even from highly active, specialized sequences. Such properties of overlapping fitness landscapes could facilitate the expansion from a weakly active, promiscuous ribozyme to an elaborated system of ribozyme-substrate pairs.

While the order in which amino acids were incorporated into the genetic code is a subject of debate, the amino acid substrates tested here include those that are generally believed to be early (L, I, V) and late (W, F, M) additions to the code^54–58^. Interestingly, the aromatic residues were generally preferred by all ribozyme families. While the original selection employed a tyrosine analog, an analogous selection using the leucine analog did not yield new ribozymes, indicating that this preference may be intrinsic. Such a preference is not surprising based on considerations for intermolecular interactions (e.g., π-π stacking) and is supported by an analysis of amino acid preferences among RNA aptamers evolved *in vitro*^72^. Thus, in a plausible scenario, self-aminoacylating RNAs that react with ‘early’ amino acid substrates would have promiscuous activity on ‘late’ substrates, allowing co-option of these ribozymes to incorporate new substrates once they become available. During code expansion, any natural selection for increased activity would also lead to increased substrate specificity, and error minimization would emerge due to the biophysical and structural preferences of the ribozymes.

These evolutionary by-products, in turn, would further improve the ability of a primitive genetic code to faithfully convert genetic information into peptide sequences with defined biophysical properties. Such emergent phenomena have been argued to be critical complements to natural selection during the origin of translation^73,74^. Like the spandrels of St. Mark’s Cathedral, architectural by-products that acquired important aesthetic value ^75^, error minimization and specificity may have originated as mechanistic by-products of how the genetic code emerged, to later become invaluable features of the modern genetic code.

## Methods

### General synthesis methods

Reagents and solvents were obtained from Sigma-Aldrich or Fisher Scientific and were used without purification, unless otherwise noted. All ^1^H NMR spectra were recorded using a Varian Unity Inova AS600 (600 MHz) with samples dissolved in DMSO-*d*6; chemical shifts δH are reported in ppm with reference to residual internal DMSO (δH = 2.50 ppm). Spectra were analyzed using MNova software.

### Preparation of biotinyl-amino acids

Biotinylation reactions were performed in 10 mL anhydrous pyridine under nitrogen. Typical reactions contained L-amino acid methyl ester hydrochloride (1 mmol), biotin (1 mmol), N-(3-dimethylaminopropyl)-N′-ethylcarbodiimide hydrochloride (EDC, 2 mmol), and 4-(dimethylamino)pyridine (0.1 mmol). The mixture was allowed to react at room temperature with stirring overnight, after which the solvent was evaporated under reduced pressure. The residue was then dissolved in dichloromethane (DCM) and washed with equal volumes of distilled water, saturated sodium bisulfate solution (twice), and saturated sodium bicarbonate solution (twice). The solution was dried with sodium sulfate, filtered, and the solvent was evaporated with reduced pressure to yield a clear, yellow solid (^1^H NMR chemical shifts reported in Table S4).

The recovered compound was dissolved by sonication in iPrOH:H_2_O (2:1 v/v) (15 mL), to which 1 mL of 3 M NaOH was added. This solution was stirred overnight at room temperature, after which the isopropyl alcohol was evaporated under reduced pressure and the product was precipitated from the remaining solution by the addition of 1 M HCl to produce a white solid. This compound was recovered by filtration, washed with water, and dried *in vacuo* (Table S4).

### Preparation of biotinyl-aminoacyl oxazolones

Oxazolone formation was performed by reacting biotinyl-amino acids (0.1 mmol) with EDC (0.12 mmol) in anhydrous DCM and stirred at 4 °C overnight. The organic phase was then washed with distilled water (twice), saturated sodium bicarbonate solution, and saturated sodium chloride solution and dried with sodium sulfate. The solution was then filtered and the solvent was evaporated under reduced pressure to yield a solid product, which was stored at −20 °C (Table S4 and Figure S9). NMR characterization was performed as described above.

Substrate solutions were prepared by weighing biotinyl-aminoacyl-oxazolone (BXO, where X = W (Trp), F (Phe), L (Leu), I (Ile), V (Val), or M (Met)) and dissolving in acetonitrile with sonication to a final concentration of 25 mM. Fresh solutions were prepared daily for each set of experiments. As a secondary means of verifying BXO concentrations in prepared solutions, a HABA biotin quantification kit (AnaSpec) was used to measure the biotin concentrations of each solution. Average measured biotin concentration and standard deviation of triplicates are shown in Table S5 (expected BXO concentration for all samples is 25 mM). While biotin quantitation measurements indicate systematically lower BXO concentrations than by weight by a factor of ~2, BXO concentrations were similar across different compounds. The low-activity background peaks also provide internal normalization to account for differences between compounds (see Results).

### Kinetic sequencing (*k*-Seq)

DNA libraries for kinetic sequencing experiments were designed as described^49^. Libraries were obtained from Integrated DNA Technologies (IDT) or Keck Biotechnology Laboratory with the sequence 5′-<u>GATAATACGACTCACTATA</u>GGGAATGGATCCACATCTACGAATTC-[central variable region, length 21]-TTCACTGCAGACTTGACGAAGCTG-3′ (nucleotides upstream of the transcription start site are underlined). The variable region was designed to contain one of the five wild-type sequences of interest (Table S1) with variability at each position corresponding to 91% wild-type base and 3% each substitution. RNA was transcribed using HiScribe T7 RNA polymerase (New England Biolabs) and purified by denaturing polyacrylamide gel electrophoresis (PAGE). Reaction pools were prepared as an equimolar mixture of each purified RNA pool and quantified by Qubit 3 Fluorometer (Invitrogen).

Kinetic sequencing experiments were performed as previously described^7,49^. Reactions were performed in 50 μL aqueous solutions containing selection buffer (100 mM HEPES, 100 mM NaCl, 100 mM KCl, 5 mM MgCl_2_, 5 mM CaCl_2_) and 5% acetonitrile at a pH between 6.9 and 7.0. Reactions contained 0.43 μM RNA and BXO at 1250, 250, 50, 10, or 2 μM. Reactions were incubated at room temperature with rotation for 90 minutes and stopped by desalting using Micro Bio-Spin Columns with Bio-Gel P-30 (Bio-Rad Laboratories). Reacted sequences were isolated with 100 μL Streptavidin MagneSphere paramagnetic beads (Promega) per sample. Beads were washed three times with PBS + 0.01% Triton X-100 and sequences were eluted into 50 μL water by heating to 70 °C for 1 minute. Samples were reverse transcribed using SuperScript III Reverse Transcriptase (Thermo Fisher Scientific). Following reverse transcription of *k*-Seq samples, qPCR reactions were performed in triplicate for each sample, including input RNA, using SsoAdvanced Universal SYBR Green Supermix (Bio-Rad Laboratories) with 2 μL of cDNA following the manufacturer’s protocol and containing 500 nM forward and reverse primers 5’GATAATACGACTCACTATAGGGAATGGATCCACATCTACGA-3’ and 5’-CAGCTTCGTCAAGTCTGCAGTGAA-3’. Serial dilutions of random library ssDNA were prepared in triplicate from 5×10^−5^ to 5×10^2^ pg/μL alongside each experiment for generating standard curves (Figure S10)^59^. Samples were analyzed using Bio-Rad CFX96 Touch system. The remaining cDNA was amplified by PCR with Phusion DNA Polymerase (Thermo Fisher Scientific) using the same forward and reverse primers as used for qPCR above. Samples were adapted for sequencing using the Nextera XT DNA Library Preparation Kit (Illumina), pooled, and sequenced by Illumina NovaSeqS4 PE150 (Novogene).

### Aminoacylation ribozyme selections

Selections for self-aminoacylating ribozymes with BFO and BLO were conducted as previously described for BYO aminoacylation^7^. Libraries were obtained from IDT with the sequence 5′-<u>GATAATACGACTCACTATA</u>GGGAATGGATCCACATCTACGAATTC-N_21_-TTCACTGCAGACTTGACGAAGCTG-3′ (T7 promoter sequence underlined), where N is an equimolar mixture of A, G, C, and T. For the first round of selection, 145 pmol of library DNA was transcribed using HiScribe T7 polymerase (New England Biolabs) and RNA was purified by gel electrophoresis. For the first round of selection, reactions contained 3.2 μM RNA and 50 μM BFO or BLO in 1 mL of selection buffer with 0.2% acetonitrile. Reactions were incubated at room temperature with rotation for 90 minutes and stopped by desalting using Micro Bio-Spin Columns with Bio-Gel P-30 (Bio-Rad Laboratories). Reacted sequences were isolated by addition of one sample volume of Streptavidin MagneSphere paramagnetic beads (Promega) per sample. Beads were washed bead buffer (PBS + 0.01% Triton X-100), 20 mM NaOH, and once more with bead buffer, then eluted by heating to 65 °C for 10 minutes in 95% formamide with 10 mM EDTA. Samples were reverse transcribed using SuperScript III Reverse Transcriptase (Thermo Fisher Scientific) and amplified with Phusion DNA Polymerase (Thermo Fisher Scientific). For subsequent rounds of selection, 7.2 pmol (round 2) or 3.6 pmol (rounds 3-5) of recovered DNA was transcribed and RNA was used at 2.2 μM in 200 μL reactions. Selections were performed for five rounds in duplicate. Samples were prepared for sequencing using the Nextera XT DNA Library Preparation Kit (Illumina), pooled, and sequenced by Illumina NextSeq 500 (Biological Nanostructures Laboratory, California NanoSystems Institute at UCSB).

### Electrophoretic mobility shift assay and determination of BFO uncatalyzed reaction rate

Gel shift assays were performed as previously described^7^. For determining the uncatalyzed reaction rate with BFO, aminoacylation reactions were performed in 50 μL selection buffer with 5% acetonitrile and contained 0.43 μM random library RNA and BFO at 1250, 250, 50, 10, or 2 μM. Reactions were incubated at room temperature for 90 minutes with rotation and stopped by desalting using Micro Bio-Spin Columns with Bio-Gel P-30 (Bio-Rad Laboratories). 95 nmol of streptavidin (New England Biolabs) was added to each sample, which were then incubated for 15 minutes with rotation at room temperature and run on an 8% polyacrylamide gel. Gel shift assays for observation of reactivity were performed with 500 μM BXO per sample unless otherwise noted.

### Computational analyses of *k*-Seq data

Sequencing reads were processed using trimmomatic SE CROP:90 to facilitate joining^76^, and then paired-end reads were joined and unique sequences were enumerated using EasyDIVER^77^. Joining was performed using the following PANDAseq^78^ flags: -a -l 1 -A pear -C completely_miss_the_point:0. These flags strip primers after assembly rather than before (-a), require sequences to have a minimum length of 1 after removing primers (-l 1), set the assembly algorithm to PEAR^79^ (-A pear), and exclude sequences with mismatches in overlapping paired-end regions (completely_miss_the_point:0). Primer sequences were extracted using CTACGAATTC as the forward primer and CTGCAGTGAA as the reverse primer.

*k*-Seq analyses were performed using the ‘k-seq’ package^49^. Briefly, the absolute quantity (ng) of a sequence in a sample was calculated as the fraction of the sequence’s read count over the total number of reads in the sample, multiplied by the mean total RNA (ng) from triplicated qPCR measurements. The input amount (ng) for a sequence was determined by the median sequence amount across 6 replicates for the unreacted pool. The fraction reacted (*F*_*S*_) was calculated as the reacted amount in the sample divided by the input amount. Sequences that contain ambiguous nucleotides (‘N’), that were not 21 nucleotides long, or that were more than two substitutions from a center sequence were excluded in downstream fitting. For each sequence, the fractions reacted in samples were fit to the pseudo-first order kinetic model 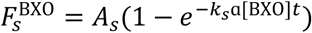, where 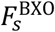 is the fraction reacted for sequence *s* with substrate BXO, *A_s_* is the maximum reaction amplitude, *k_s_* is the rate constant, and [BXO] is the initial concentration of BXO. *α* is the coefficient accounting for the hydrolysis of substrate BXO during the reaction time (*t* = 90 min), and a fixed value (0.479, measured for BYO^7^) was used for all substrates. Note that the effect of α on estimated *k_s_* cancels out when calculating the catalytic enhancement ratio *r_s_*. To quantify the estimation uncertainty of kinetic model parameters (*k_s_*, *A_s_*) for each sequence, samples (fractions reacted) were bootstrapped (resampling with replacement to the original size) for 1000 times and each bootstrapped sample set was fit into the model for *k*_*S*_ and *A*_*S*_. Statistics (e.g., median, standard deviation, 2.5-percentile, 97.5 percentile) were calculated from bootstrapped results. The median of product *k_s_A_s_* was used to represent the activity of each sequence.

### Background reaction rate estimation

Histograms (100 bins) of log_10_-transformed *kA* values for sequences from all families were fit to a bimodal Gaussian distribution (Figure S4 and Table S2). The mean of the low-activity peak (*μ_1_*) was used as the estimated uncatalyzed rate (*k_0_A_0_*) and the standard deviation of the fit (*σ_1_*) was used to inform the choice of catalytic enhancement threshold. Additionally, the uncatalyzed reaction rate was calculated for BFO by gel shift assay as described previously for BYO^7^ (see above).

### Clustering analysis of sequences from selections

Sequences were clustered into families based on sequence similarity, using a custom Python script (see Data Availability). The script *ClusterBOSS.py* uses the enumerated read output files generated from the EasyDIVER package^77^. In general, first, all sequences were sorted according to their read count values. Then, the most abundant sequence was chosen as a candidate ‘center’ sequence to start a family, as long as its read count value was at least 10 (*c_min_* = 10). The Levenshtein edit distance (number of substitutions, insertions, or deletions) from this candidate sequence to every other sequence in the distribution was computed (no restriction on minimum number of counts; *a*_*min*_ = 1). If the distance was less than a cutoff (*d_cutoff_* = 3 mutations from the center sequence), the sequence was considered to be part of the same family as the initially chosen center sequence. No restriction was applied to the number of sequences required to define a family (*n_min_* = 1), which includes the center sequence and any sequences found to cluster with it. Once assigned to a family, sequences were not allowed to be clustered into another family. To find the rest of the family clusters, we followed the same procedure until all sequences had been explored.

### Promiscuity indices

Promiscuity indices were calculated using the calculator available at http://hetaira.herokuapp.com/. Due to the single-turnover nature of the aminoacylation ribozymes studied here, promiscuity indices are calculated using catalytic enhancement values instead of the catalytic efficiency as originally described by Nath and Atkins^61^.

## Data availability

Data from high-throughput sequencing and *k*-Seq analysis (Figures 2-7) will be available at the Dryad Digital Repository (https://doi.org/10.25349/D92C9C).

Private link for peer review: https://datadryad.org/stash/share/OSZh1pcJLVz_cJabbozcA8x68wDJET-f485TeN_54_s.

## Code availability

Scripts not reported elsewhere are available at https://github.com/ichen-lab-ucsb/ClusterBOSS (ClusterBOSS: Cluster Based On Sequence Similarity) and https://github.com/ichen-lab-ucsb/WFLIVM_k-Seq (scripts used to generate figures in this manuscript).

## Acknowledgements

The authors thank John Sutherland for support for organic synthesis and discussion of exaptation, Jen Smith for advice on sequencing, Huan Peng, Yei-Chen Lai, Josh Kenchel, and Abe Pressman for technical advice, Robert Pascal for advice on organic synthesis, and William Atkins and Abhinav Nath for advice on the promiscuity index. The authors acknowledge the use of the Biological Nanostructures Laboratory within the California NanoSystems Institute, supported by the University of California, Santa Barbara and the University of California, Office of the President. Funding from the Simons Foundation Collaboration on the Origin of Life (290356FY18), NASA (NNX16AJ32G), National Institute of General Medical Sciences (DP2GM123457), NSF (1935372) and the Camille Dreyfus Teacher-Scholar Program is acknowledged.

## Author contributions

E.J. and I.A.C. designed the project; E.J. conducted experiments and analyzed data; E.J. and Z.L. synthesized oxazolones; Y.S. analyzed *k*-Seq data; C.B. wrote the clustering script; E.J. and I.A.C. interpreted data; E.J. and I.A.C. wrote the manuscript with input from all authors.

## Notes

https://doi.org/10.25349/D92C9C

https://github.com/ichen-lab-ucsb/ClusterBOSS

https://github.com/ichen-lab-ucsb/WFLIVM_k-Seq

### Competing Interest Statement

The authors have declared no competing interest.

