## Supplementary Material for "Error minimization and specificity could emerge in a genetic code as by-products of prebiotic evolution"

**Table S1.** Wild-type sequences for ribozyme families used in this study.

| Family | Wild-type sequence for 21-nt selected region |
|---|---|
| 1A.1 | CTACTTCAAACAATCGGTCTG |
| 1B.1 | CCACACTTCAAGCAATCGGTC |
| 2.1 | ATTACCCTGGTCATCGAGTGA |
| 2.2 | ATTCACCTAGGTCATCGGGTG |
| 3.1 | AAGTTTGCTAATAGTCGCAAG |

**Figure S1.** Duplicate selections (A and B) for aminoacylating ribozymes with BFO and BLO result in convergence on the same primary families identified previously under selection with BYO[1]. Note that only Motif 2 emerges in substantial fraction during selection with BLO, while the BFO selection yields families from multiple Motifs.

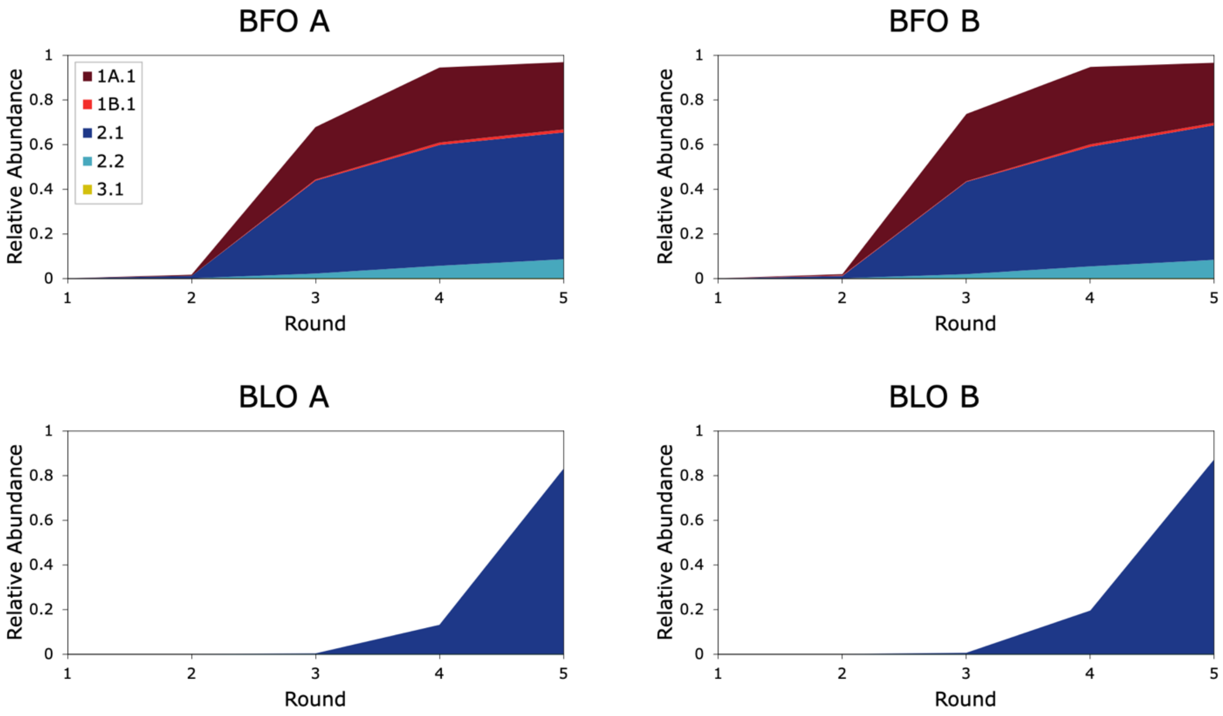

**Figure S2.** Precision of *k*-Seq estimates of $k_sA_s$. Bootstrapping (N=1000) was used to estimate 95% confidence intervals (95% CI range, i.e. 97.5%-2.5%) and medians as previously described[2]. Confidence intervals were normalized to the medians estimated from bootstrapping. It can be seen that normalized confidence intervals are generally within one order of magnitude.

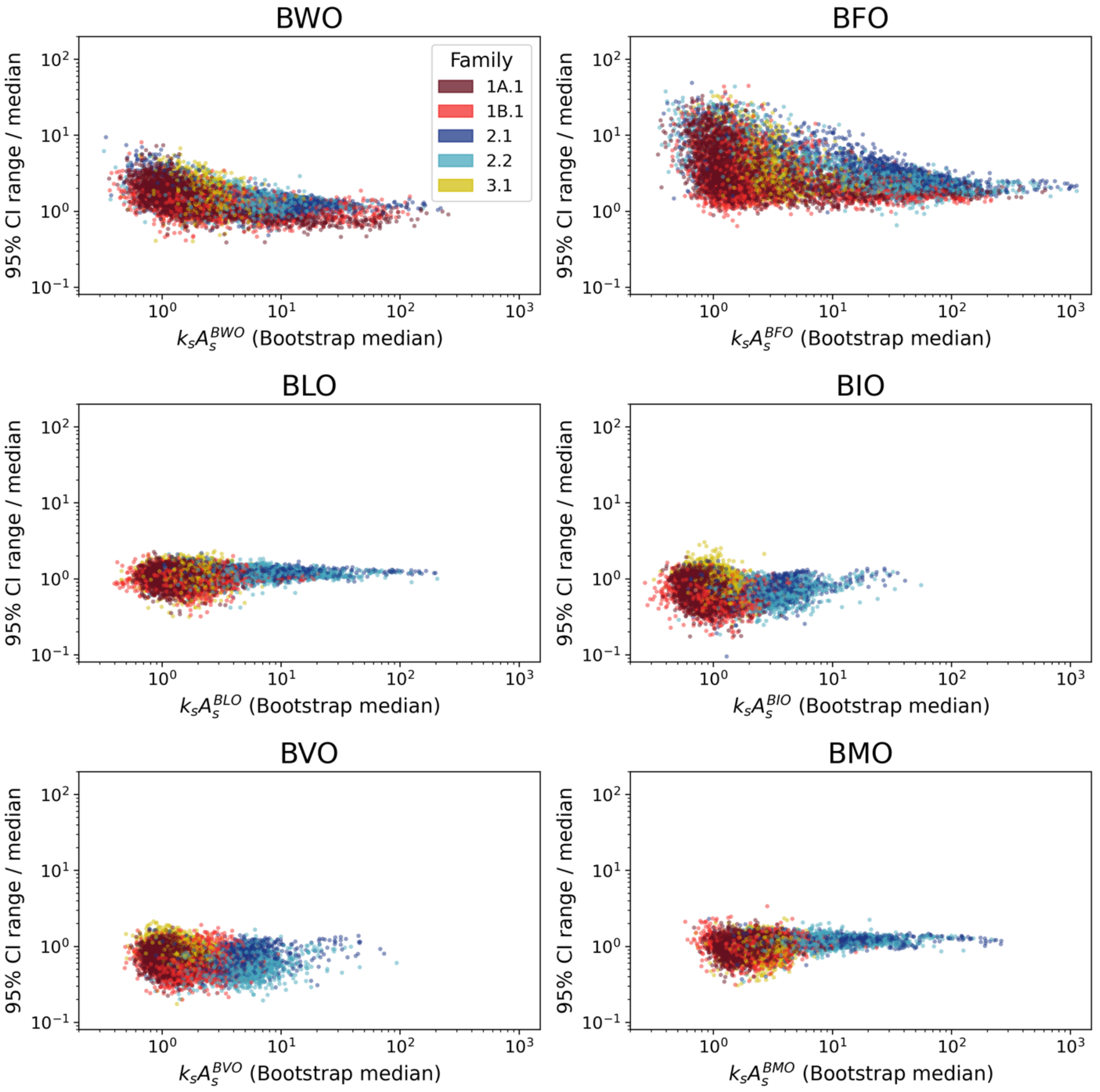

**Figure S3.** Histograms of ribozyme $k_sA_s$ values with each substrate, for each family (see legend).

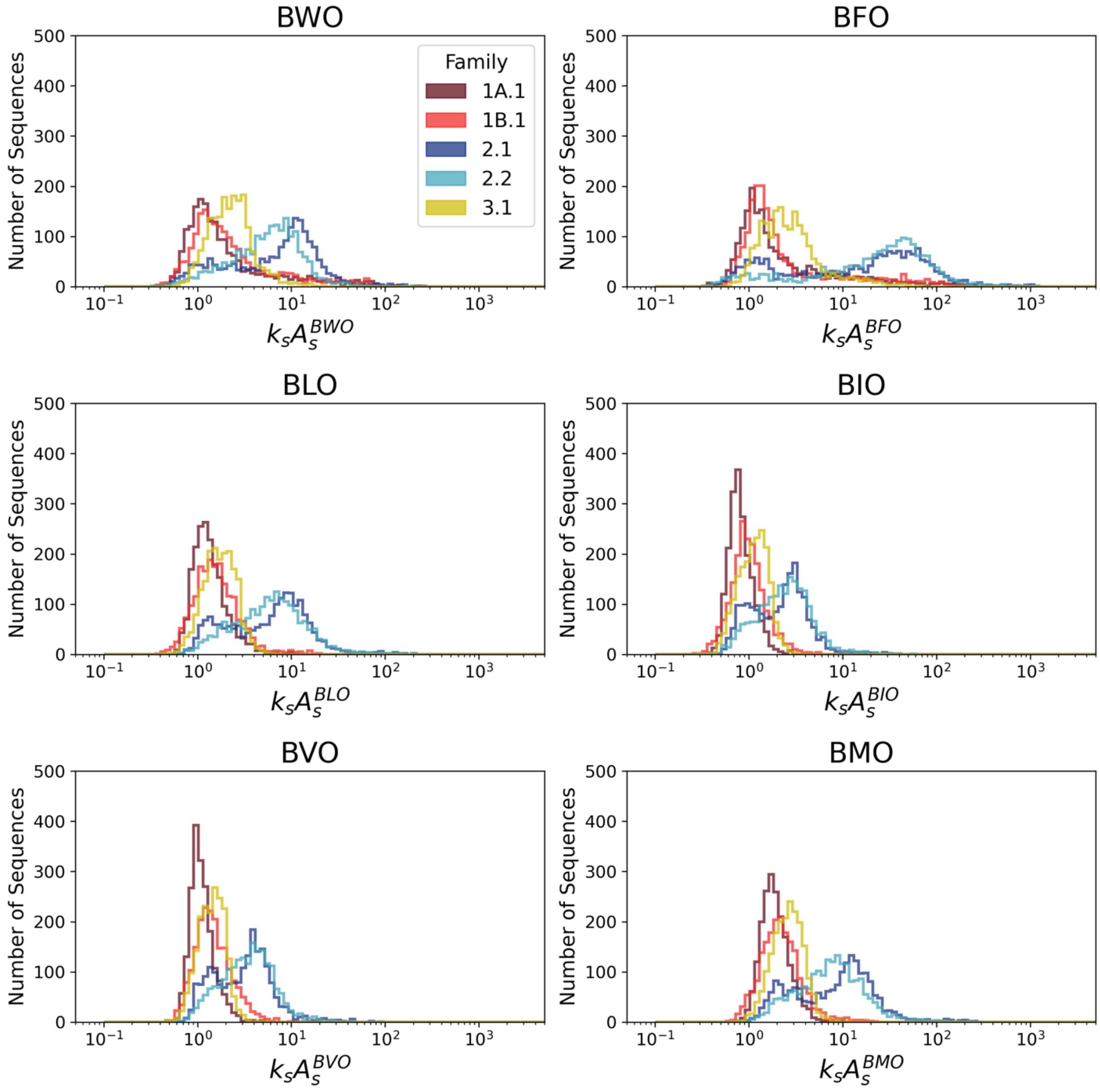

**Figure S4.** Frequency distribution (histogram) of $\log_{10}$-transformed $k_sA_s$ for the ribozyme variants reacted with each substrate. The frequency distribution of ribozymes has been previously found to be log-normal[3]. Bimodal Gaussian fits (black lines) were used to characterize the low-activity peak using Equation 1 below. The centers of the low-activity peaks ($\mu_1$) and their standard deviations ($\sigma_1$) are given in Table S2.

**Equation 1**: $y = a_1 e^{-\left(\frac{x-\mu_1}{2\sigma_1}\right)^2} + a_2 e^{-\left(\frac{x-\mu_2}{2\sigma_2}\right)^2}$

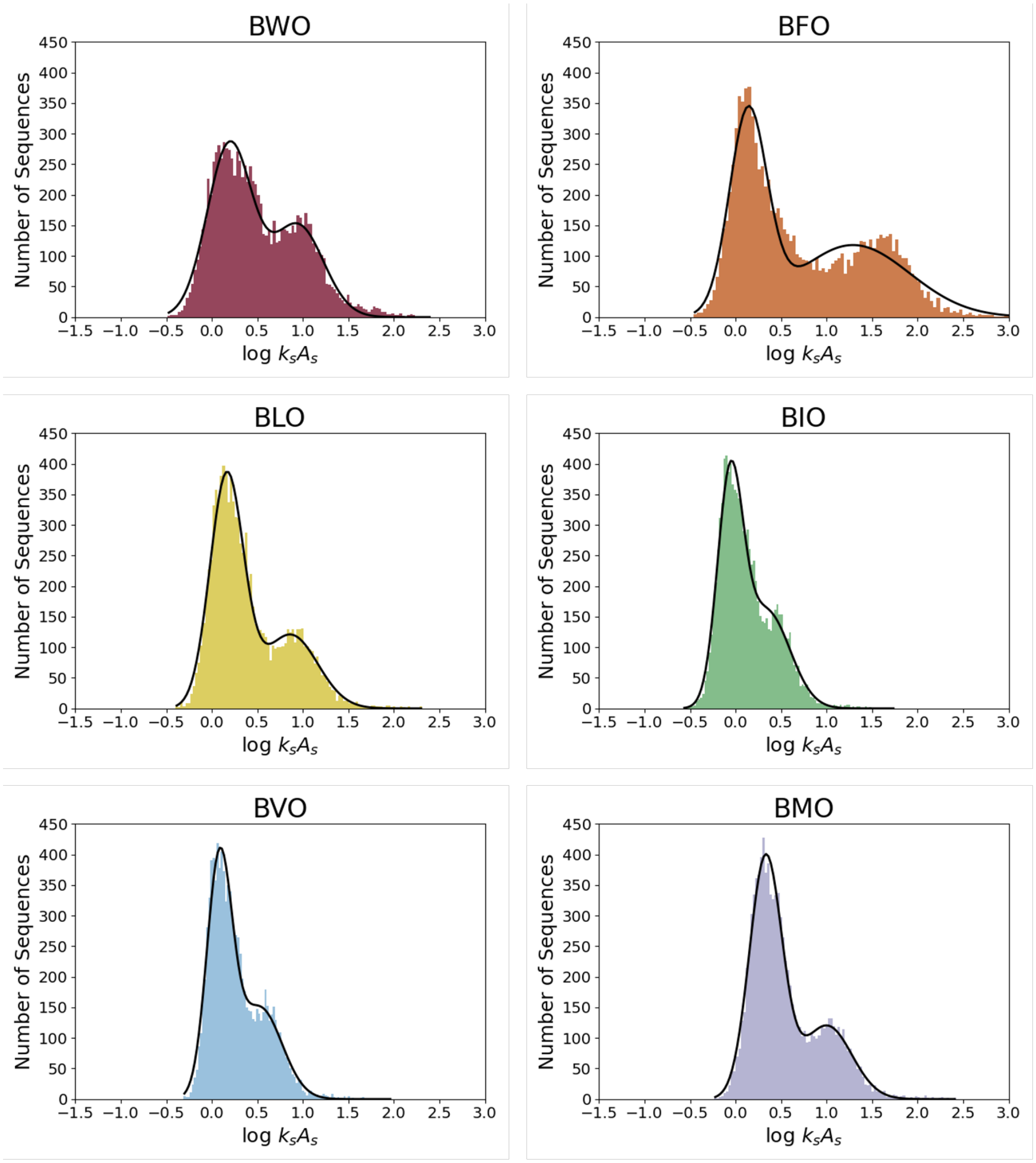

**Table S2.** Characterization of the background peaks. The center of the low-activity peak (Figure S3) corresponded to the background rate of BFO measured by gel shift. $10^{\mu_1}$ of the low-activity peak was used as the presumed background rate ($k_0A_0$) for each substrate.

| **Substrate** | $\mu_1$ | $\sigma_1$ | **2.5%** | **97.5%** | **$10^{\mu_1}$ ($M^{-1}min^{-1}$)** | **95% range ($M^{-1}min^{-1}$)** |
|---:|---:|---:|---:|---:|---:|---:|
| BWO | 0.196 | 0.247 | -0.289 | 0.680 | 1.57 | 0.51-4.79 |
| BFO | 0.138 | 0.205 | -0.265 | 0.541 | 1.37 | 0.54-3.47 |
| BLO | 0.164 | 0.185 | -0.200 | 0.527 | 1.46 | 0.63-3.36 |
| BIO | -0.060 | 0.141 | -0.337 | 0.217 | 0.87 | 0.46-1.65 |
| BVO | 0.083 | 0.140 | -0.191 | 0.357 | 1.21 | 0.64-2.28 |
| BMO | 0.331 | 0.180 | -0.021 | 0.684 | 2.14 | 0.95-4.83 |

**Figure S5.** Histograms of catalytic enhancement for all families with each substrate. The same data are represented in Figure 2 (but presented by family instead of by substrate).

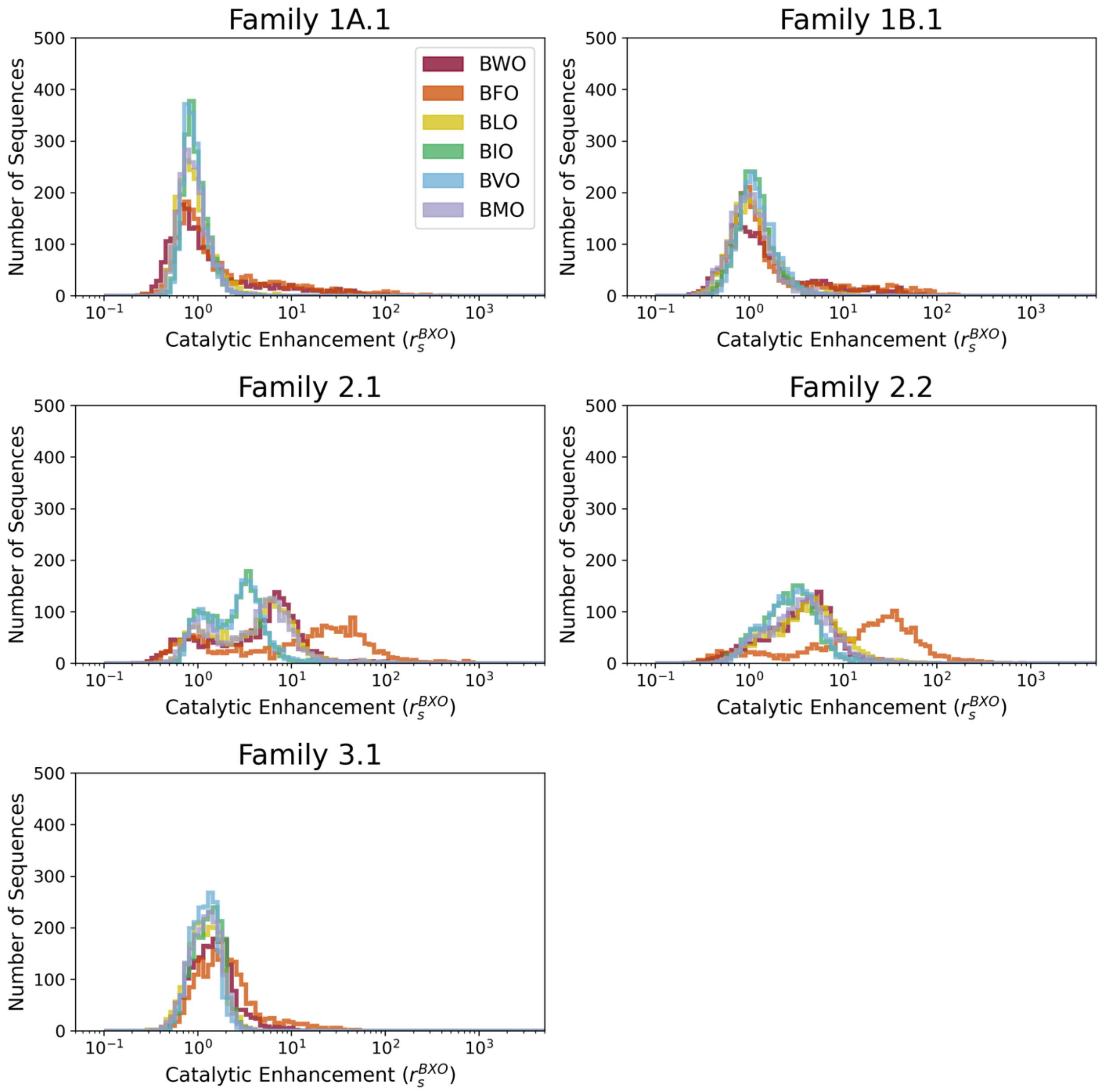

**Figure S6.** Pairwise comparisons of catalytic enhancement values for each substrate for Families A) 1A.1, B) 1B.1, C) 2.1, D) 2.2, and E) 3.1. Wild-type sequences are shown in red, single-mutants are shown in blue, and double-mutants are shown in yellow. Dashed gray line indicates line of identity. Black lines indicate linear regression fits used to calculate $R^2$ values and slopes in Figure 3. The same data are also plotted in Figure 3, but here the families are plotted separately, with mutant order and error bars (95% confidence interval) indicated. 95% confidence intervals of $r_s$ were calculated from confidence intervals of $k_sA_s$ with normalization by the constant $k_0A_0$.

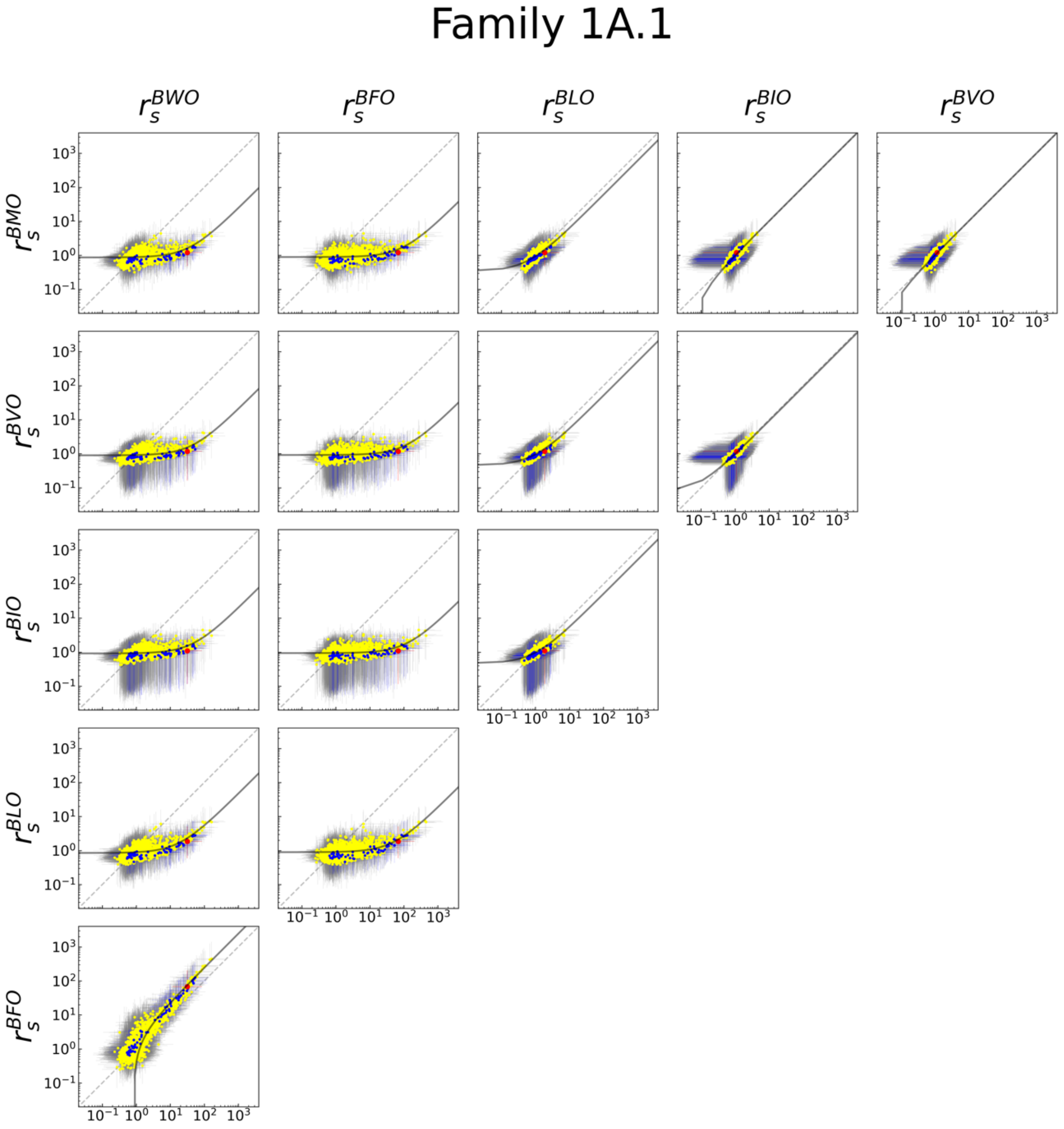

Figure S6, continued.

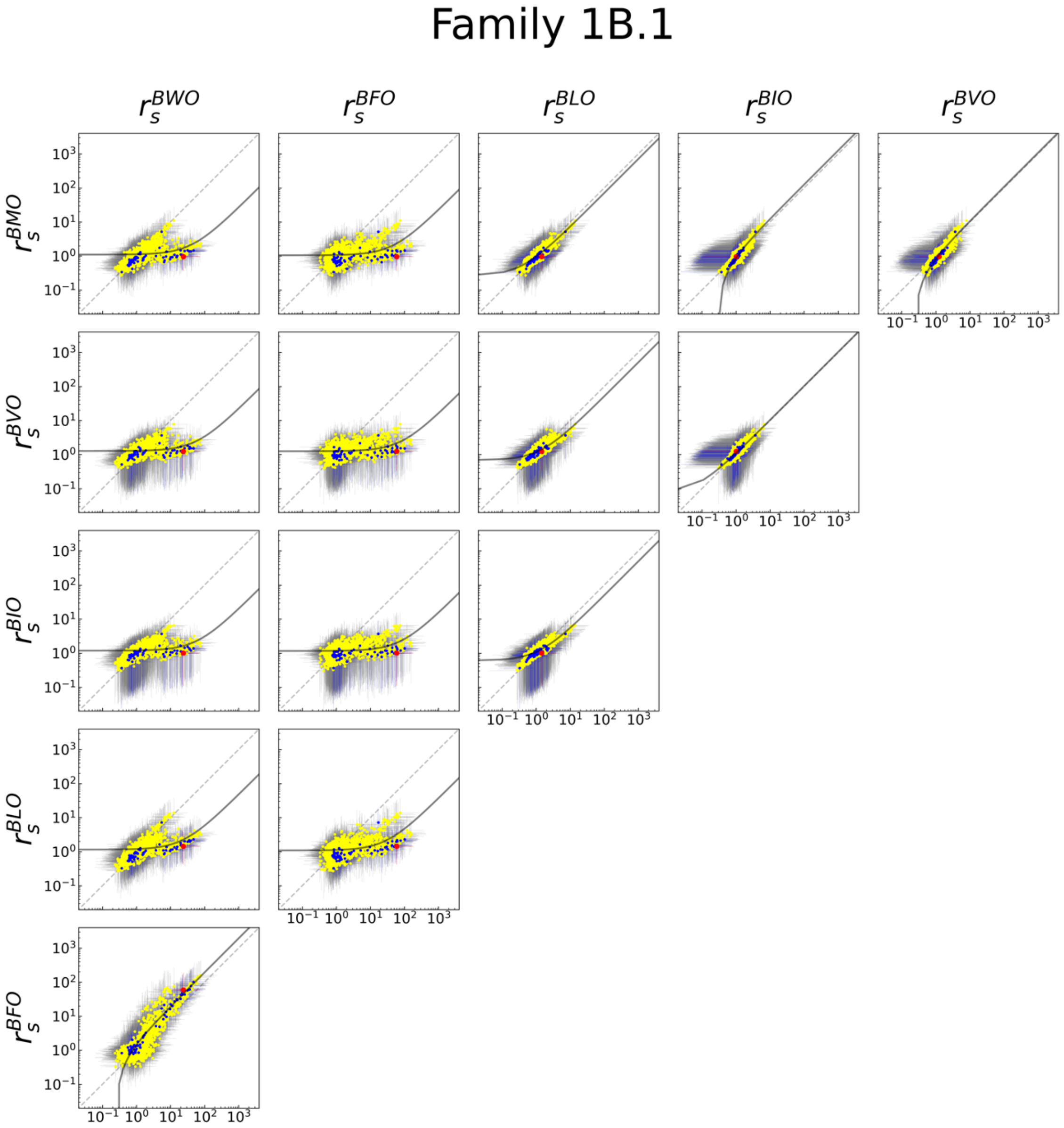

Figure S6, continued.

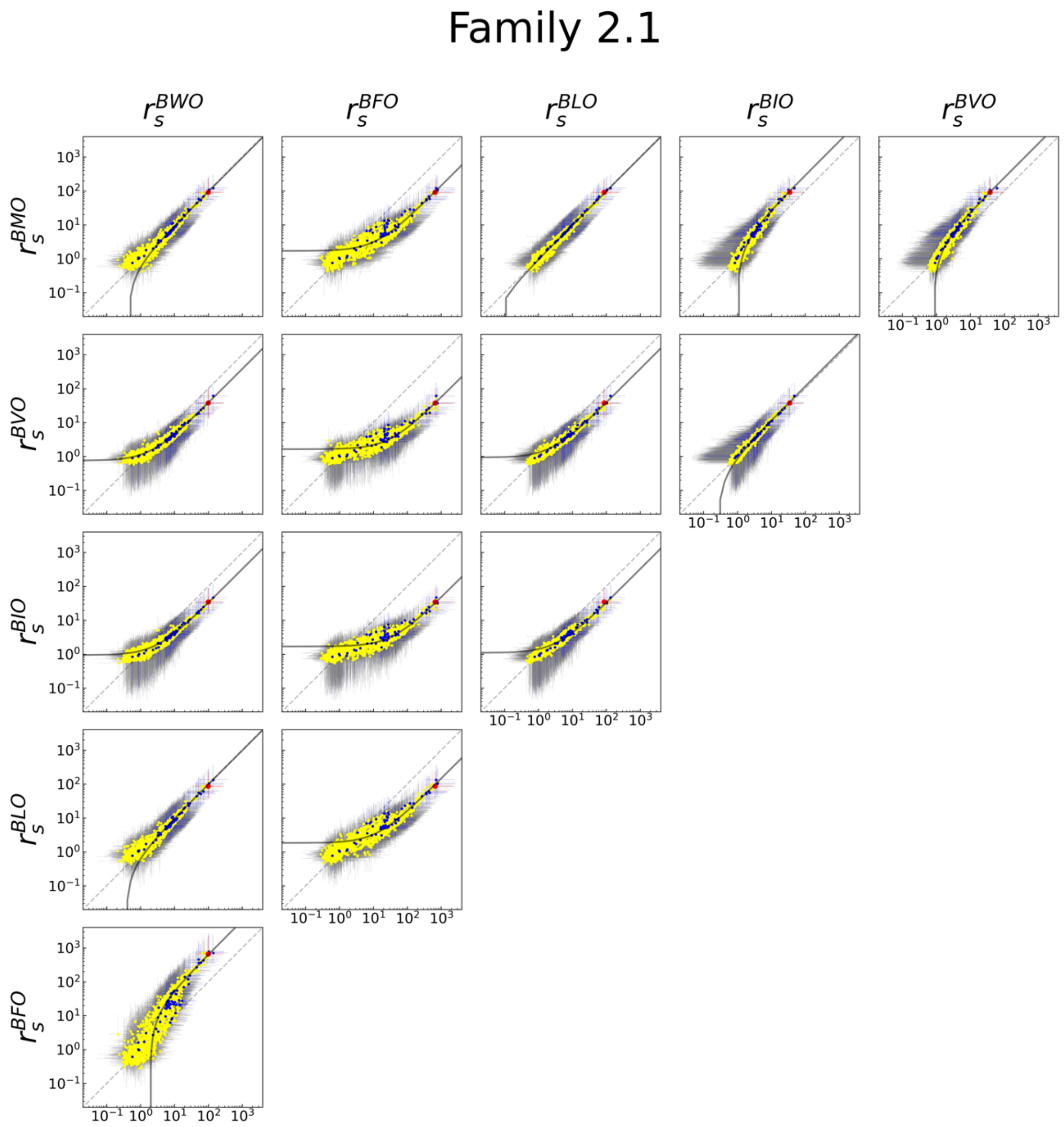

Figure S6, continued.

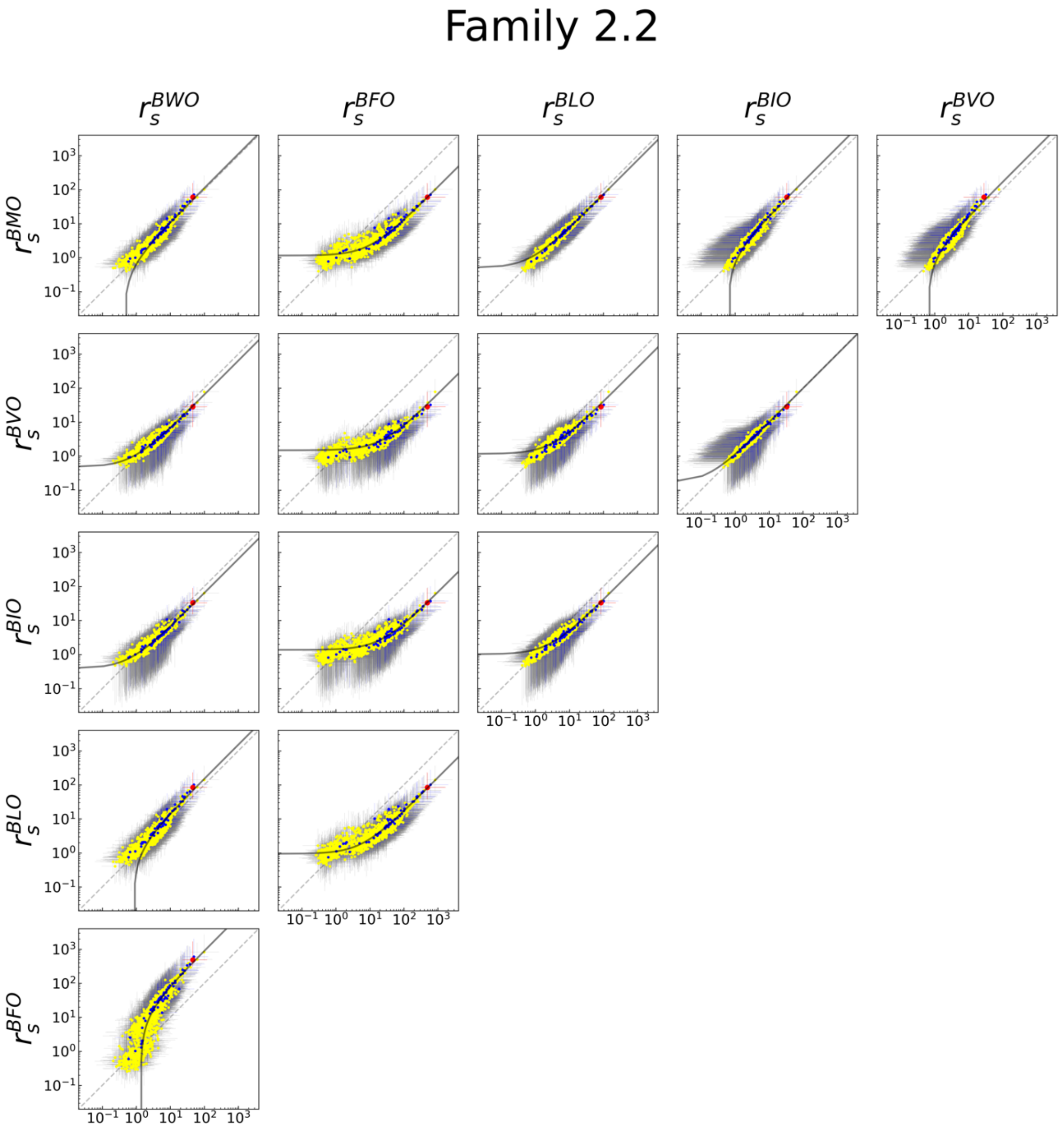

Figure S6, continued.

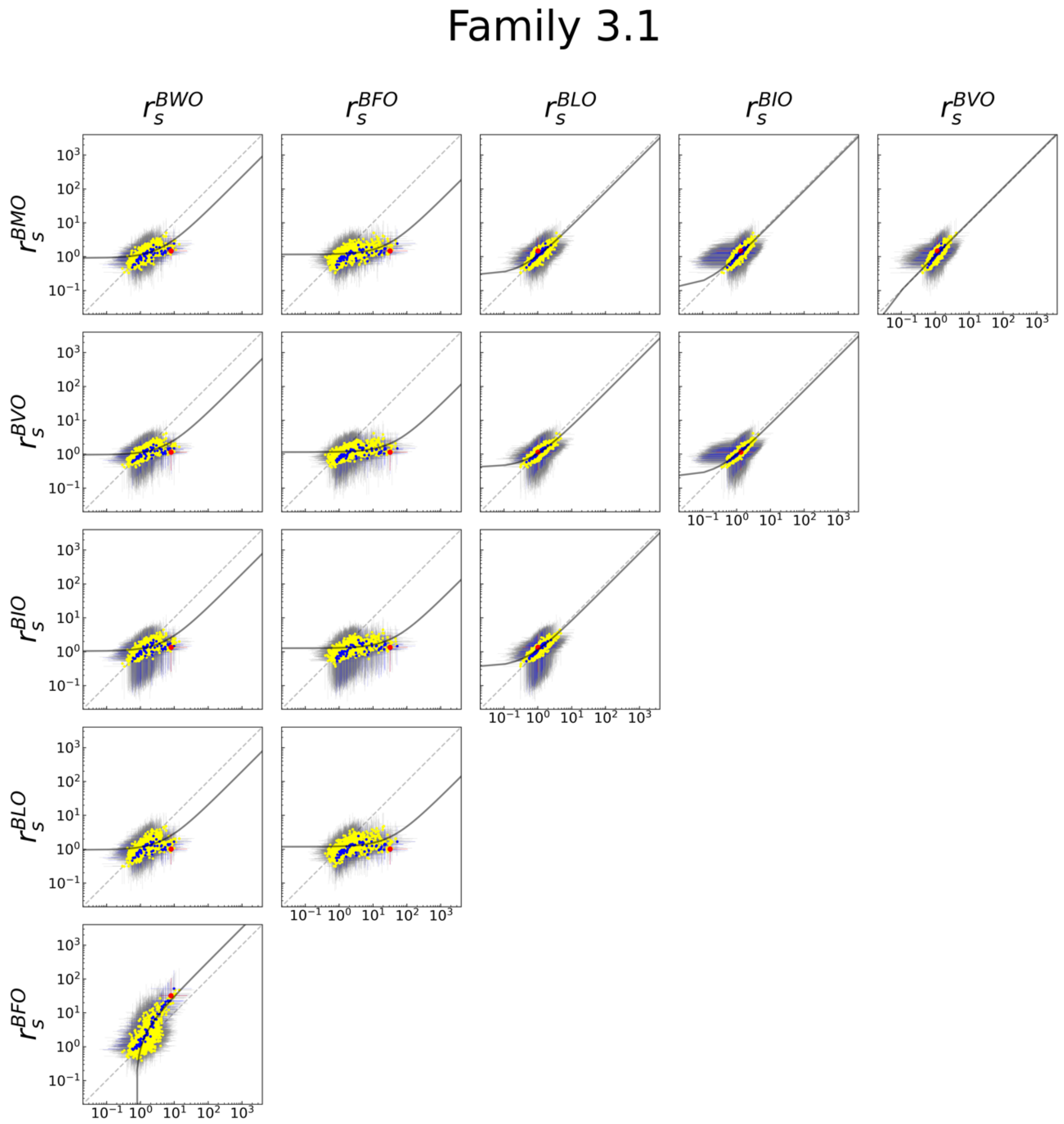

**Figure S7.** Relationship of activity to promiscuity index $I_s$, separated by family. These plots show the same data as Figure 5, with families plotted separately.

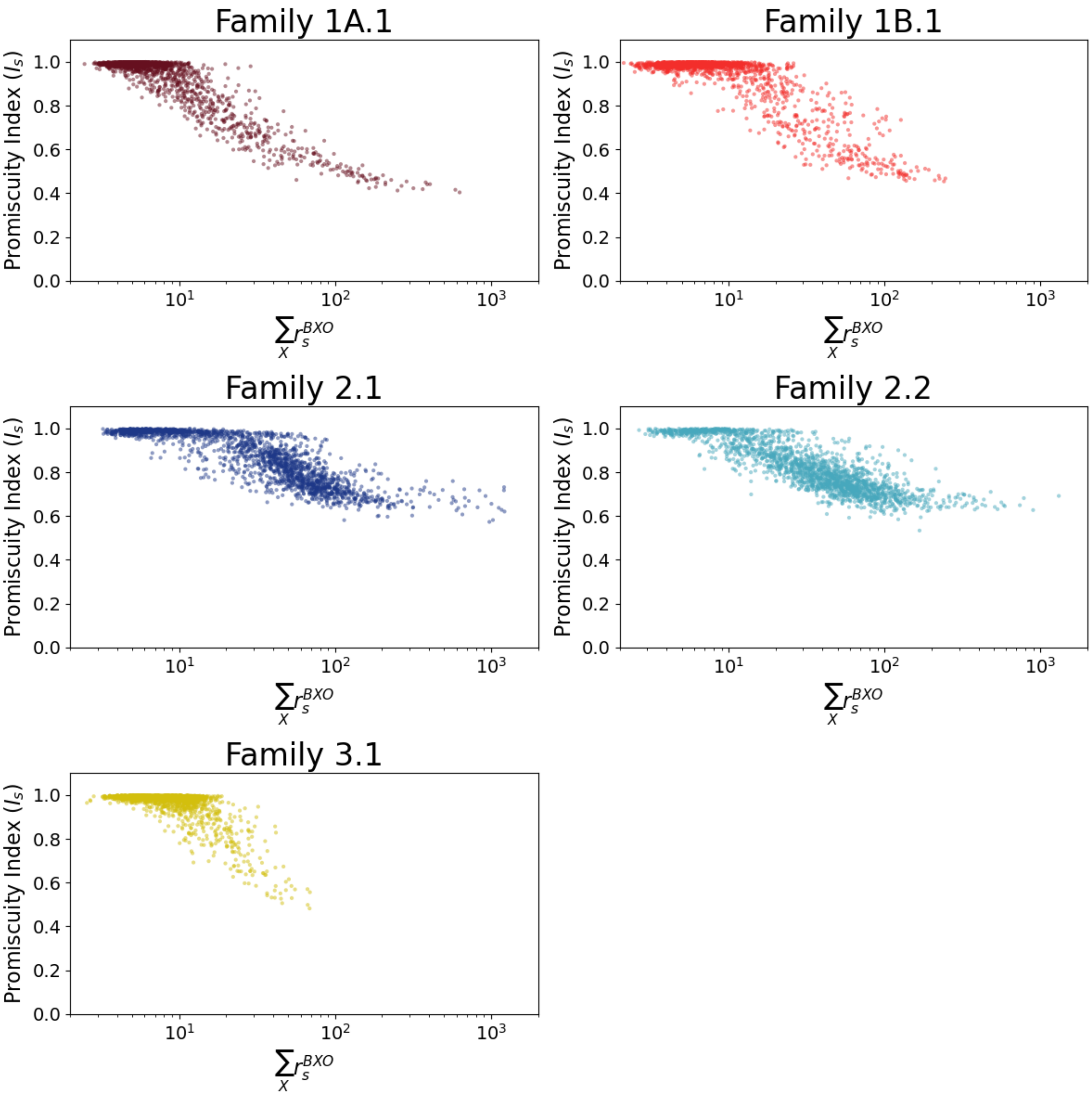

**Figure S8.** Observed preference for aromatic substrates, as determined by the ratio of the sum of activity on BWO and BFO to the sum of activity on all tested substrates (BXO) (aromatic preference ratio). Increasing preference can be observed for increasing activity.

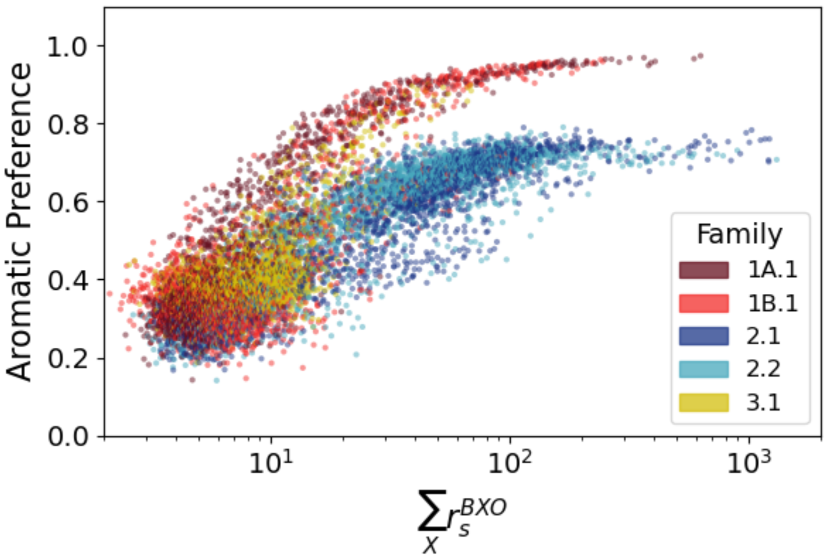

**Table S3.** Catalytic enhancement values for all sequences shown in Figure 7. See Data Availability for raw data and confidence intervals. The (X,Y) coordinates of each sequence in each family plot of Figure 7 is given, where the coordinate (0,0) is the middle of the plot.

| Family 1A.1 Sequences | $r_s^{BWO}$ | $r_s^{BFO}$ | $r_s^{BLO}$ | $r_s^{BIO}$ | $r_s^{BVO}$ | $r_s^{BMO}$ | X | Y |
|---|---|---|---|---|---|---|---|---|
| CTACTTCAAACAATCGGTCTG | 31.4 | 67.9 | 1.9 | 1.1 | 1.2 | 1.2 | 0.173 | 0.114 |
| CTGCTTCAAACAATCGGCCTG | 162.4 | 449.5 | 6.9 | 3.1 | 3.3 | 3.7 | -0.709 | 0.428 |
| CCACTTCAAACAATCGGCCTG | 144.9 | 425.3 | 7.5 | 4.3 | 4 | 4.7 | -0.548 | -0.402 |
| CAACTTCAAACAATCGGCCTG | 106.7 | 282.5 | 6.8 | 3.1 | 2.9 | 4.1 | -0.292 | -0.557 |
| CGACTTCAAACAATCGGCCTG | 95.7 | 276.7 | 6.7 | 3.3 | 4.1 | 3.6 | -0.007 | -0.442 |
| GTACTTCAAACAATCGGCCTG | 89.6 | 271.6 | 7 | 4.5 | 4.2 | 4.1 | 0.108 | 0.531 |
| CCGCTTCAAACAATCGGTCTG | 95.9 | 257.1 | 4.8 | 1.8 | 2.4 | 2.7 | -0.666 | 0.110 |
| GTGCTTCAAACAATCGGTCTG | 79.6 | 195.2 | 4.9 | 2.6 | 2.4 | 2.7 | -0.007 | 0.987 |
| CTGCTTCAAACAATCGGTCTG | 53.5 | 125.3 | 2.8 | 1.3 | 1.4 | 1.7 | -0.306 | 0.541 |
| CCACTTCAAACAATCGGTCTG | 49.3 | 113.7 | 2.8 | 1.5 | 1.6 | 1.8 | -0.158 | -0.224 |
| CGACTTCAAACAATCGGTCTG | 44.4 | 101.6 | 2.7 | 1.5 | 1.5 | 1.7 | 0.428 | -0.270 |
| CAACTTCAAACAATCGGTCTG | 44.8 | 97.7 | 2.6 | 1.6 | 1.5 | 1.7 | 0.220 | -0.425 |
| CTACTTCAAACAATCGGCCTG | 48.1 | 85.3 | 2.9 | 1.7 | 1.8 | 1.9 | -0.262 | 0.037 |
| GTACTTCAAACAATCGGTCTG | 30.5 | 51.3 | 2.1 | 1.5 | 1.4 | 1.5 | 0.486 | 0.550 |
| CCACTTCAAACAATCGGGCTG | 27.2 | 27.3 | 2.6 | 2.1 | 3 | 2.1 | -0.294 | -0.825 |
| CAACTTCAAACAATCGGGCTG | 20.1 | 23.8 | 2.6 | 2.2 | 2.3 | 1.9 | 0.040 | -0.954 |
| GTACTTCAAACAATCGGGCTG | 14.6 | 21.1 | 2.7 | 3 | 2.8 | 2.5 | 0.458 | 0.938 |
| CGACTTCAAACAATCGGGCTG | 12.7 | 10.4 | 2.3 | 2.4 | 2.7 | 2.1 | 0.335 | -0.796 |
| GTACTTCAAACAATCGGTGTG | 10.2 | 10.4 | 3.4 | 2.1 | 2 | 2.8 | 1.000 | 0.661 |
| **Family 1B.1 Sequences** | $r_s^{BWO}$ | $r_s^{BFO}$ | $r_s^{BLO}$ | $r_s^{BIO}$ | $r_s^{BVO}$ | $r_s^{BMO}$ | X | Y |
| CCACACTTCAAGCAATCGGTC | 23.9 | 59.3 | 1.4 | 1 | 1.2 | 1 | -0.071 | 0.027 |
| CCGCGCTTCAAGCAATCGGTC | 77.2 | 157.8 | 3.2 | 2 | 2.7 | 2.4 | -0.935 | -0.320 |
| CCGCACTTCAAACAATCGGTC | 74.3 | 156.9 | 3.4 | 1.6 | 2 | 1.9 | -0.381 | -1.000 |
| CCGCACTTCAACCAATCGGTC | 67.5 | 147.6 | 3.4 | 2.1 | 2.4 | 2.1 | 0.040 | -0.987 |
| CCCCGCTTCAAGCAATCGGTC | 70.4 | 141.6 | 3.7 | 2 | 2.7 | 2.1 | -0.956 | -0.022 |
| CCCCACTTCAAACAATCGGTC | 61.2 | 139.8 | 2.8 | 1.9 | 2.1 | 2 | -0.605 | -0.725 |
| GCACGCTTCAAGCAATCGGTC | 60.1 | 120.2 | 3.6 | 2.6 | 2.7 | 2.6 | -0.826 | 0.695 |
| CCCCACTTCAACCAATCGGTC | 53.6 | 101.2 | 3 | 2.4 | 1.8 | 1.9 | -0.200 | -0.807 |
| CCACACTTCAAACAATCGGTC | 47 | 101.9 | 2.3 | 1.4 | 1.5 | 1.5 | -0.105 | -0.483 |
| CCCCACTTCAAGCAATCGGTG | 12.5 | 75.4 | 13.9 | 6.6 | 7.8 | 11 | 0.266 | -0.075 |
| CCACGCTTCAAGCAATCGGTC | 39.9 | 76.9 | 2.1 | 1.2 | 1.5 | 1.4 | -0.648 | 0.203 |
| CCACACTTCAAGCATTCGGTG | 7.9 | 68.5 | 11 | 5.5 | 6 | 8.9 | 0.870 | -0.116 |
| CCGCACTTCAAGCAATCGGTC | 36.5 | 61.6 | 2 | 1.3 | 1.7 | 1.4 | -0.365 | -0.471 |
| CCACATTTCAAGCAATCGGTG | 9.4 | 59.1 | 12.9 | 6.2 | 5.9 | 8.9 | 0.993 | 0.092 |
| CCACACTTCAACCAATCGGTC | 35.2 | 60.5 | 2.2 | 1.4 | 1.5 | 1.4 | 0.212 | -0.520 |
| CCACAATTCAAGCAATCGGTG | 8.8 | 60.3 | 12.3 | 6.4 | 6 | 8.1 | 0.996 | 0.377 |
| ACACACTTCAAGCAATCGGTG | 9.8 | 56.7 | 10.5 | 5.9 | 5.8 | 7.8 | 0.363 | 0.746 |
| CCCCACTTCAAGCAATCGGTC | 31.5 | 57.5 | 2 | 1.5 | 1.8 | 1.5 | -0.373 | -0.219 |
| GCACACTTCAAGCAATCGGTG | 10.1 | 49.1 | 10.3 | 6.2 | 5.9 | 8.1 | 0.106 | 0.820 |
| CCACACTTCAAGAAATCGGTG | 8.4 | 55.3 | 10.6 | 4.8 | 3.5 | 6.5 | 0.884 | 0.625 |
| CCACACTTCAAGGAATCGGTG | 6.6 | 52.1 | 10.5 | 4.9 | 4.6 | 7.2 | 0.673 | 0.837 |
| GCACACTTCAAGCAATCGGTC | 19.8 | 32.3 | 1.8 | 1.7 | 1.7 | 1.5 | -0.350 | 0.555 |
| ACACACTTCAAGCAATCGGTC | 13.1 | 27.1 | 1.3 | 1.2 | 1.4 | 1.1 | -0.067 | 0.463 |
| CCACACTTCAAGCAATCGGTG | 5.4 | 17.1 | 7.2 | 3.7 | 3.7 | 5.2 | 0.479 | 0.304 |
| **Family 2.1 Sequences** | $r_s^{BWO}$ | $r_s^{BFO}$ | $r_s^{BLO}$ | $r_s^{BIO}$ | $r_s^{BVO}$ | $r_s^{BMO}$ | X | Y |

| | | | | | | | | |
|---|---|---|---|---|---|---|---|---|
| ATTACCCTGGTCATCGAGTGA | 99.1 | 656.9 | 86.9 | 34.6 | 37.9 | 91.3 | 0.111 | 0.128 |
| GTTACCCTGGTCATCGAGTGT | 102.2 | 825.3 | 113.5 | 31.5 | 37.1 | 111.2 | 0.595 | -0.711 |
| ATTACCCTGGTCATCGGGTGA | 140 | 711.2 | 132.7 | 47 | 60.9 | 121.9 | -0.408 | -0.106 |
| ATTACCCTGGTCATCGGGTGT | 132.7 | 721.9 | 136.1 | 40.7 | 56.5 | 123.1 | -0.069 | -0.549 |
| ATTACCCTGGTCATCGAGTGT | 103.8 | 785.1 | 104.6 | 31.7 | 37.4 | 107.7 | 0.484 | -0.241 |
| GTTACCCTGGTCATCGAGTGA | 101.3 | 747.3 | 100.3 | 35.2 | 38.3 | 99 | 0.124 | -0.431 |
| ATGGCCCTGGTCATCGAGTGA | 76.5 | 728.9 | 84.6 | 27.1 | 29.1 | 82.9 | 0.106 | 1.000 |
| ATGACCCTGGTCATCGGGTGA | 94.9 | 571 | 99 | 31.9 | 35.1 | 90.6 | -0.682 | 0.346 |
| GTTACCCTGGTCATCGGGTGA | 90.7 | 503.5 | 92.1 | 30.9 | 36.7 | 89.7 | -0.481 | -0.598 |
| ATGACCCTGGTCATCGAGTGA | 51.6 | 451.3 | 55.8 | 18.6 | 19.3 | 58.9 | -0.208 | 0.582 |
| ATTGCCCTGGTCATCGAGTGA | 60 | 367.3 | 51.2 | 20.6 | 23.7 | 53.1 | 0.429 | 0.580 |
| **Family 2.2 Sequences** | $r_s^{BWO}$ | $r_s^{BFO}$ | $r_s^{BLO}$ | $r_s^{BIO}$ | $r_s^{BVO}$ | $r_s^{BMO}$ | **X** | **Y** |
| ATTCACCTAGGTCATCGGGTG | 45.6 | 485.8 | 83.7 | 33.2 | 28.4 | 60.1 | -0.145 | 0.021 |
| ATTCCCCTAGGTCATCGCGTG | 99.5 | 826.3 | 140.9 | 64.1 | 77.6 | 101.6 | 0.577 | -0.703 |
| ATTCACCTAGGTCATCGGGTA | 49.5 | 605.2 | 100.6 | 37.2 | 32 | 71.8 | -0.501 | 0.387 |
| ATTCCCCTAAGTCATCGGGTG | 62.2 | 498.1 | 85.8 | 38.6 | 37.9 | 59.1 | 0.767 | 0.047 |
| ATTCGCCTAGGTCATCGGGTA | 35.3 | 390.7 | 68.8 | 25.1 | 23.4 | 46.9 | 0.007 | 0.543 |
| GTTCACCTAGGTCATCGGGTA | 33.1 | 380.8 | 68.3 | 24.6 | 22.6 | 50.6 | -0.620 | 0.877 |
| ATTCGCCTAGGTCATCGGGTG | 37.3 | 364.4 | 63.8 | 26.2 | 23.1 | 45.8 | 0.258 | 0.048 |
| ATTGACCTAGGTCATCGGGTA | 31.5 | 355.2 | 59.5 | 20.4 | 16.7 | 47.1 | -1.000 | 0.290 |
| ATTGACCTAGGTCATCGGGTG | 31.1 | 337.2 | 58.8 | 20.6 | 17.4 | 43.8 | -0.676 | -0.052 |
| ATTCCCCTAGGTCATCCGGTG | 39.5 | 295.2 | 51.5 | 27.6 | 26.9 | 38.5 | -0.060 | -0.804 |
| ATTCCCCTAGGTCATCGGGTG | 23.2 | 196.4 | 32.9 | 14.9 | 14.9 | 24.4 | 0.281 | -0.274 |
| GTTCACCTAGGTCATCGGGTG | 20.1 | 179.9 | 32.5 | 13.7 | 12.5 | 25.5 | -0.279 | 0.559 |
| ATTCGCCTAGGTCATCGCGTG | 17.6 | 125.6 | 23.4 | 10 | 10.9 | 19.2 | 0.683 | -0.333 |
| ATTCGCCTAAGTCATCGGGTG | 11.7 | 117.3 | 15.6 | 7.8 | 7.9 | 12.9 | 0.709 | 0.404 |
| ATTCACCTAGGTCATCGCGTG | 11.5 | 100.5 | 17.4 | 7.1 | 7.3 | 13 | 0.186 | -0.465 |
| ATTCGCCTAGGTCATCCGGTG | 10.3 | 78.7 | 10.9 | 6.5 | 6.7 | 8 | -0.086 | -0.466 |
| ATTCACCTAAGTCATCGGGTG | 8.9 | 74.2 | 12.8 | 5.8 | 5.6 | 9.7 | 0.338 | 0.382 |
| ATTCACCTAGGTCATCCGGTG | 6.3 | 44.9 | 8.4 | 4.7 | 4.7 | 6.3 | -0.438 | -0.462 |
| **Family 3.1 Sequences** | $r_s^{BWO}$ | $r_s^{BFO}$ | $r_s^{BLO}$ | $r_s^{BIO}$ | $r_s^{BVO}$ | $r_s^{BMO}$ | **X** | **Y** |
| AAGTTTGCTAATAGTCGCAAG | 8 | 32 | 1 | 1.3 | 1.1 | 1.5 | -0.312 | 0.346 |
| GAGTCTGCTAATAGTCGCAAG | 14 | 46.9 | 2.1 | 2 | 1.6 | 2.3 | -0.270 | -0.184 |
| AAGTCTGCTAATAGTCGCAAG | 9.7 | 51.8 | 1.7 | 1.5 | 1.5 | 2.4 | 0.278 | -0.085 |
| AAGTCTGCTAATAGTCGTAAG | 9.9 | 49.6 | 1.7 | 1.6 | 1.5 | 2.5 | -0.296 | -0.671 |
| AAGTCCGCTAATAGTCGCAAG | 10.7 | 46.3 | 2 | 2.4 | 2.1 | 3 | 0.941 | 0.090 |
| AAGCCTGCTAATAGTCGCAAG | 11.1 | 37.3 | 1.6 | 1.9 | 1.8 | 1.6 | 0.837 | -0.530 |
| AGGTCTGCTAATAGTCGCAAG | 10 | 37 | 1.1 | 1.4 | 1.2 | 2.1 | 0.433 | 0.620 |
| TAGTCTGCTAATAGTCGCAAG | 9.9 | 32.7 | 1.8 | 2 | 1.8 | 2.2 | 0.202 | -0.666 |
| AAGTTTGCTAATAGTCGTAAG | 8.2 | 33 | 0.9 | 1.2 | 1 | 1.3 | -0.854 | -0.258 |
| AAGTTTGCTAATAGTCGCGCG | 5.7 | 20.9 | 4.8 | 3.5 | 2.9 | 4 | -0.854 | -0.671 |
| AAGCTTGCTAATAGTCGCGAG | 3.8 | 26.9 | 2.7 | 2.7 | 1.9 | 3.4 | 0.488 | 0.620 |
| AAGTTTGCTAATAGTCGCGGG | 4.8 | 19.6 | 4.5 | 4.4 | 3.8 | 4.1 | -0.003 | -0.881 |
| GAGTTTGCTAATAGTCGCAAG | 7.4 | 22.7 | 1.4 | 1.7 | 1.5 | 1.8 | -0.850 | 0.338 |
| AACTTTGCTAATAGTCGCGAG | 3.1 | 19.5 | 3.7 | 3.5 | 2.8 | 3 | -0.850 | 0.314 |
| AGGTTTGCTAATAGTCGCAAG | 6.6 | 22 | 1.4 | 1.6 | 1.4 | 1.8 | -0.109 | 1.000 |
| AAGTTTGCTAATAGTCGCCGG | 5.8 | 10.5 | 3.5 | 3.1 | 4.1 | 5 | 0.855 | -1.061 |
| AACTTTGCTAAGAGTCGCAAG | 5 | 12.8 | 4.1 | 2.8 | 2.6 | 3.9 | -0.255 | 0.789 |
| AACTTCGCTAATAGTCGCAAG | 3.1 | 14.5 | 3.4 | 3 | 3 | 3.5 | 0.818 | -0.066 |

**Table S4.** $^{1}$H NMR spectra for synthesized compounds.

| Compound | 1H NMR Spectrum |
|---|---|
| Biotinyl-tryptophan methyl ester | $^{1}$H NMR (600 MHz, DMSO-d6) δ = 10.82 (s, 1H), 8.19 (d, J = 7.6 Hz, 1H), 7.47 (d, J = 7.9 Hz, 1H), 7.31 (d, J = 8.1 Hz, 1H), 7.11 (d, J = 2.3 Hz, 1H), 7.04 (dd, J = 11.1, 4.0 Hz, 1H), 6.96 (t, J = 7.5 Hz, 1H), 6.35 (s, 1H), 6.32 (s, 1H), 4.48 (dd, J = 13.8, 8.2 Hz, 1H), 4.30 – 4.25 (m, 1H), 4.09 – 4.04 (m, 1H), 3.56 (s, 3H), 3.11 (dd, J = 14.6, 5.5 Hz, 1H), 3.01 (m, 2H), 2.80 (dd, J = 12.4, 5.1 Hz, 1H), 2.55 (d, J = 12.4 Hz, 1H), 2.12 – 2.00 (m, 2H), 1.60 – 1.19 (m, 6H). |
| Biotinyl-tryptophan | $^{1}$H NMR (600 MHz, DMSO-d6) δ = 12.53 (s, 1H), 10.79 (s, 1H), 8.03 (d, J = 7.9 Hz, 1H), 7.50 (d, J = 7.8 Hz, 1H), 7.31 (d, J = 8.1 Hz, 1H), 7.10 (d, J = 2.2 Hz, 1H), 7.04 (t, J = 7.5 Hz, 1H), 6.96 (t, J = 7.1 Hz, 1H), 6.35 (s, 1H), 6.32 (s, 1H), 4.45 (td, J = 8.5, 5.1 Hz, 1H), 4.30 – 4.25 (m, 1H), 4.08 – 4.03 (m, 1H), 3.13 (dd, J = 14.6, 5.0 Hz, 1H), 3.03 – 2.93 (m, 2H), 2.79 (dd, J = 12.4, 5.1 Hz, 1H), 2.55 (d, J = 12.4 Hz, 1H), 2.11 – 1.99 (m, 2H), 1.60 – 1.13 (m, 6H). |
| Biotinyl-tryptophan oxazolone (BWO) | $^{1}$H NMR (600 MHz, DMSO-d6) δ = 10.86 (s, 1H), 7.50 (d, J = 7.9 Hz, 1H), 7.29 (dd, J = 8.1, 2.6 Hz, 1H), 7.06 (t, J = 2.6 Hz, 1H), 7.03 (dd, J = 11.1, 4.0 Hz, 1H), 6.94 (dd, J = 11.0, 3.9 Hz, 1H), 6.37 (s, 1H), 6.33 (s, 1H), 4.67 (d, J = 4.9 Hz, 1H), 4.33 – 4.24 (m, 1H), 4.09 – 4.04 (m, 1H), 3.23 (dd, J = 14.5, 4.6 Hz, 1H), 3.13 (ddd, J = 14.8, 6.0, 1.9 Hz, 1H), 2.96 (ddd, J = 17.9, 11.6, 6.6 Hz, 1H), 2.81 (dt, J = 12.4, 4.9 Hz, 1H), 2.56 (d, J = 12.4 Hz, 1H), 2.31 – 2.18 (m, 2H), 1.56 – 1.07 (m, 6H). |
| Biotinyl-phenylalanine methyl ester | $^{1}$H NMR (600 MHz, DMSO-d6) δ = 8.24 (d, J = 7.8 Hz, 1H), 7.30 – 7.14 (m, 5H), 6.37 (s, 1H), 6.33 (s, 1H), 4.44 (td, J = 9.3, 5.5 Hz, 1H), 4.32 – 4.25 (m, 1H), 4.13 – 4.04 (m, 1H), 3.57 (d, J = 8.5 Hz, 3H), 3.08 – 2.96 (m, 2H), 2.85 (dd, J = 13.7, 9.7 Hz, 1H), 2.81 (dd, J = 12.4, 5.1 Hz, 1H), 2.56 (d, J = 12.4 Hz, 1H), 2.09 – 1.98 (m, 2H), 1.60 – 1.12 (m, 6H). |
| Biotinyl-phenylalanine | $^{1}$H NMR (600 MHz, DMSO-d6) δ = 12.57 (s, 1H), 8.03 (d, J = 8.1 Hz, 1H), 7.25 – 7.09 (m, 5H), 6.32 (s, 1H), 6.28 (s, 1H), 4.39 – 4.32 (m, 1H), 4.26 – 4.22 (m, 1H), 4.06 – 4.01 (m, 1H), 3.03 – 2.94 (m, 2H), 2.81 – 2.73 (m, 2H), 2.52 (d, J = 12.4 Hz, 1H), 2.03 – 1.92 (m, 2H), 1.57 – 1.07 (m, 6H). |
| Biotinyl-phenylalanine oxazolone (BFO) | $^{1}$H NMR (600 MHz, DMSO-d6) δ = 7.31 – 7.12 (m, 5H), 6.41 (s, 1H), 6.33 (s, 1H), 4.72 – 4.66 (m, 1H), 4.32 – 4.26 (m, 1H), 4.14 – 4.07 (m, 1H), 3.12 (dd, J = 14.0, 5.1 Hz, 1H), 3.07 – 3.02 (m, 1H), 2.96 (dd, J = 14.0, 6.8 Hz, 1H), 2.81 (ddd, J = 12.4, 5.1, 2.2 Hz, 1H), 2.57 (d, J = 12.4 Hz, 1H), 2.39 – 2.25 (m, 2H), 1.62 – 1.16 (m, 6H). |
| Biotinyl-leucine methyl ester | $^{1}$H NMR (600 MHz, DMSO-d6) δ = 8.13 (d, J = 7.7 Hz, 1H), 6.37 (s, 1H), 6.33 (s, 1H), 4.31 – 4.27 (m, 1H), 4.26 – |

| | |
|---|---|
| | 4.21 (m, 1H), 4.13 – 4.08 (m, 1H), 3.59 (s, 3H), 3.10 – 3.03 (m, 1H), 2.80 (dd, J = 12.4, 5.1 Hz, 1H), 2.56 (d, J = 12.4 Hz, 1H), 2.10 (t, J = 7.3 Hz, 2H), 1.67 – 1.19 (m, 9H), 0.84 (dd, J = 32.3, 6.6 Hz, 6H). |
| Biotinyl-leucine | $^{1}$H NMR (600 MHz, DMSO-d6) δ = 12.37 (s, 1H), 7.94 (d, J = 8.0 Hz, 1H), 6.32 (s, 1H), 6.28 (s, 1H), 4.26 – 4.21 (m, 1H), 4.14 (dd, J = 14.0, 9.0 Hz, 1H), 4.06 (d, J = 6.0 Hz, 1H), 3.02 (t, J = 9.3 Hz, 1H), 2.76 (dd, J = 12.4, 5.1 Hz, 1H), 2.51 (d, J = 12.4 Hz, 1H), 2.05 (t, J = 7.2 Hz, 2H), 1.62 – 1.21 (m, 9H), 0.80 (dd, J = 32.1, 6.5 Hz, 6H). |
| Biotinyl-leucine oxazolone (BLO) | $^{1}$H NMR (600 MHz, DMSO-d6) δ = 6.41 (s, 1H), 6.33 (s, 1H), 4.35 (dd, J = 8.1, 6.5 Hz, 1H), 4.31 – 4.26 (m, 1H), 4.14 – 4.09 (m, 1H), 3.12 – 3.06 (m, 1H), 2.81 (dd, J = 12.4, 5.1 Hz, 1H), 2.56 (d, J = 12.4 Hz, 1H), 2.44 (tt, J = 10.8, 5.4 Hz, 2H), 1.86 – 1.33 (m, 9H), 0.89 (dd, J = 14.2, 6.7 Hz, 6H). |
| Biotinyl-isoleucine methyl ester | $^{1}$H NMR (600 MHz, DMSO-d6) δ = 8.00 (d, J = 8.0 Hz, 1H), 6.32 (s, 1H), 6.27 (s, 1H), 4.26 – 4.20 (m, 1H), 4.13 (t, J = 7.4 Hz, 1H), 4.06 (d, J = 2.6 Hz, 1H), 3.54 (s, 3H), 3.02 (dd, J = 6.4, 3.9 Hz, 1H), 2.75 (dd, J = 12.4, 5.0 Hz, 1H), 2.50 (d, J = 12.4 Hz, 1H), 2.15 – 1.99 (m, 2H), 1.73 – 1.05 (m, 9H), 0.76 (dd, J = 7.0, 5.3 Hz, 6H). |
| Biotinyl-isoleucine | $^{1}$H NMR (600 MHz, DMSO-d6) δ = 12.46 (s, 1H), 7.90 (d, J = 8.4 Hz, 1H), 6.37 (s, 1H), 6.32 (s, 1H), 4.31 – 4.26 (m, 1H), 4.15 (dd, J = 8.3, 6.3 Hz, 1H), 4.13 – 4.08 (m, 1H), 3.07 (dt, J = 8.6, 6.1 Hz, 1H), 2.80 (dd, J = 12.4, 5.1 Hz, 1H), 2.56 (d, J = 12.4 Hz, 1H), 2.19 – 2.07 (m, 2H), 1.78 – 1.11 (m, 9H), 0.86 – 0.79 (m, 6H). |
| Biotinyl-isoleucine oxazolone (BIO) | $^{1}$H NMR (600 MHz, DMSO-d6) δ = 6.41 (s, 1H), 6.33 (s, 1H), 4.34 (ddt, J = 32.9, 4.1, 1.9 Hz, 1H), 4.30 – 4.26 (m, 1H), 4.14 – 4.10 (m, 1H), 3.13 – 3.05 (m, 1H), 2.81 (dd, J = 12.4, 5.1 Hz, 1H), 2.56 (d, J = 12.4 Hz, 1H), 2.46 (dd, J = 7.3, 1.9 Hz, 2H), 1.92 – 1.14 (m, 9H), 0.94 – 0.69 (m, 6H). |
| Biotinyl-valine methyl ester | $^{1}$H NMR (600 MHz, DMSO-d6) δ = 7.99 (d, J = 8.0 Hz, 1H), 6.33 (s, 1H), 6.28 (s, 1H), 4.27 – 4.21 (m, 1H), 4.12 – 4.02 (m, 2H), 3.56 (s, 3H), 3.07 – 2.99 (m, 1H), 2.76 (dd, J = 12.3, 5.0 Hz, 1H), 2.51 (d, J = 12.5 Hz, 1H), 2.15 – 2.04 (m, 2H), 1.94 (dt, J = 13.1, 6.5 Hz, 1H), 1.59 – 1.18 (m, 6H), 0.80 (dd, J = 12.1, 6.8 Hz, 6H). |
| Biotinyl-valine | $^{1}$H NMR (600 MHz, DMSO-d6) δ = 12.47 (s, 1H), 7.88 (d, J = 8.5 Hz, 1H), 6.37 (s, 1H), 6.32 (s, 1H), 4.31 – 4.25 (m, 1H), 4.11 (m, 2H), 3.07 (dt, J = 8.6, 6.0 Hz, 1H), 2.80 (dd, J = 12.4, 5.1 Hz, 1H), 2.55 (d, J = 12.4 Hz, 1H), 2.21 – 2.08 (m, 2H), 2.01 (dq, J = 13.4, 6.7 Hz, 1H), 1.66 – 1.22 (m, 6H), 0.85 (dd, J = 6.8, 1.9 Hz, 6H). |
| Biotinyl-valine oxazolone (BVO) | $^{1}$H NMR (600 MHz, DMSO-d6) δ = 6.41 (s, 1H), 6.33 (s, 1H), 4.31 – 4.24 (m, 2H), 4.14 – 4.09 (m, 1H), 3.13 – 3.05 (m, 1H), 2.81 (dd, J = 12.4, 5.1 Hz, 1H), 2.56 (d, J = 12.4 |

| | |
|---|---|
| | Hz, 1H), 2.46 (dd, J = 8.9, 6.9 Hz, 2H), 2.15 – 2.06 (m, 1H), 1.67 – 1.32 (m, 6H), 1.00 – 0.79 (m, 6H). |
| Biotinyl-methionine methyl ester | $^1$H NMR (600 MHz, DMSO-d6) δ = 8.18 (d, J = 7.5 Hz, 1H), 6.37 (s, 1H), 6.33 (s, 1H), 4.37 – 4.31 (m, 1H), 4.31 – 4.26 (m, 1H), 4.15 – 4.08 (m, 1H), 3.60 (s, 3H), 3.10 – 3.04 (m, 1H), 2.80 (dd, J = 12.4, 5.1 Hz, 1H), 2.56 (d, J = 12.4 Hz, 1H), 2.53 – 2.38 (m, 2H), 2.10 (t, J = 7.4 Hz, 2H), 2.02 (s, 3H), 1.95 – 1.78 (m, 2H), 1.66 – 1.20 (m, 6H). |
| Biotinyl-methionine | $^1$H NMR (600 MHz, DMSO-d6) δ = 12.53 (s, 1H), 8.05 (d, J = 7.8 Hz, 1H), 6.37 (s, 1H), 6.33 (s, 1H), 4.32 – 4.24 (m, 2H), 4.14 – 4.08 (m, 1H), 3.12 – 3.03 (m, 1H), 2.80 (dd, J = 12.4, 5.1 Hz, 1H), 2.56 (d, J = 12.4 Hz, 1H), 2.51 – 2.38 (m, 2H), 2.10 (t, J = 7.3 Hz, 2H), 2.02 (s, 3H), 1.97 – 1.75 (m, 2H), 1.67 – 1.21 (m, 6H). |
| Biotinyl-methionine oxazolone (BMO) | $^1$H NMR (600 MHz, DMSO-d6) δ = 6.41 (s, 1H), 6.33 (s, 1H), 4.45 (td, J = 5.7, 2.0 Hz, 1H), 4.29 (dd, J = 7.6, 5.2 Hz, 1H), 4.15 – 4.09 (m, 1H), 3.09 (dd, J = 12.6, 6.4 Hz, 1H), 2.81 (dd, J = 12.4, 5.1 Hz, 1H), 2.59 – 2.38 (m, 5H), 2.08 – 1.83 (m, 5H), 1.63 – 1.37 (m, 6H). |

**Figure S9.** $^{1}$H NMR spectra for BXO compounds shown in Figure 1A.

Biotinyl-tryptophan oxazolone (BWO)

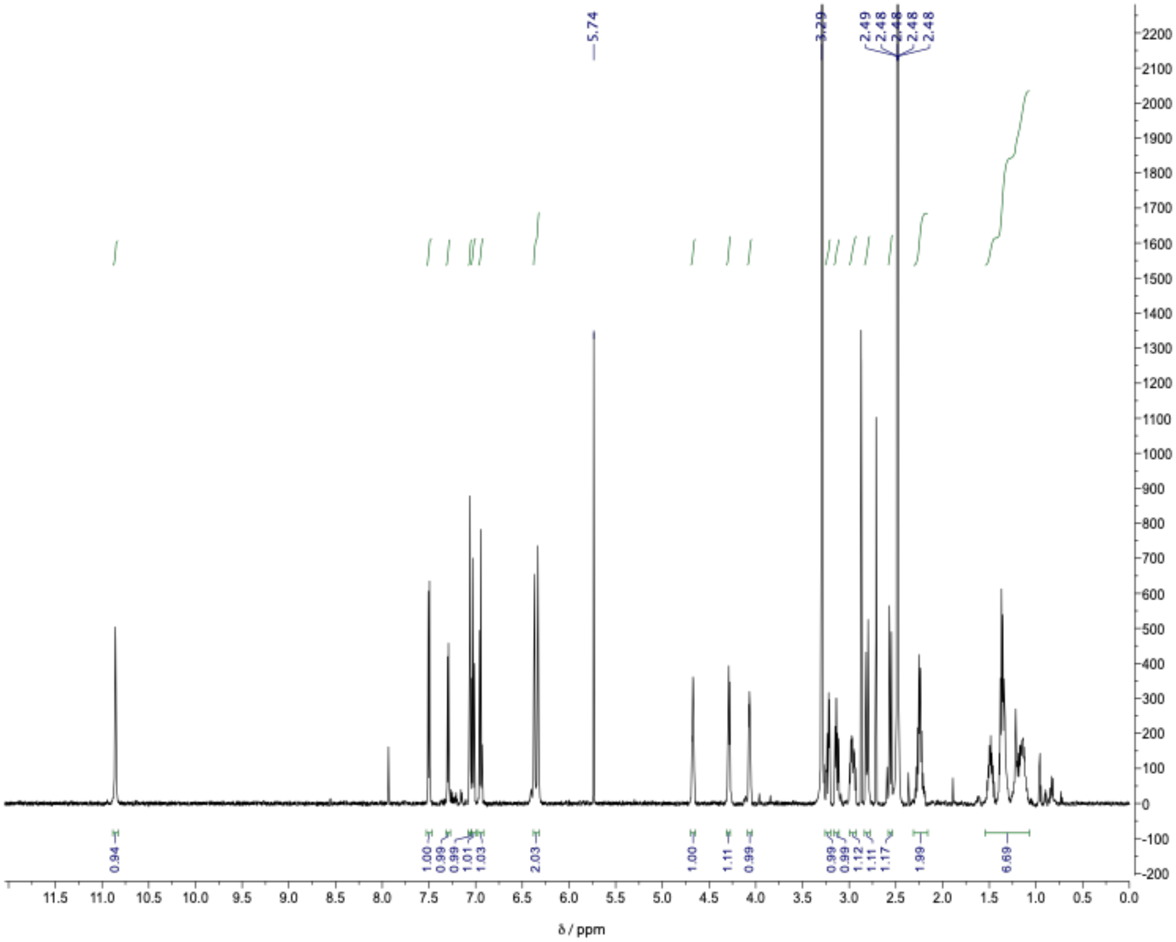

Figure S9, continued.
Biotinyl-phenylalanine oxazolone (BFO)

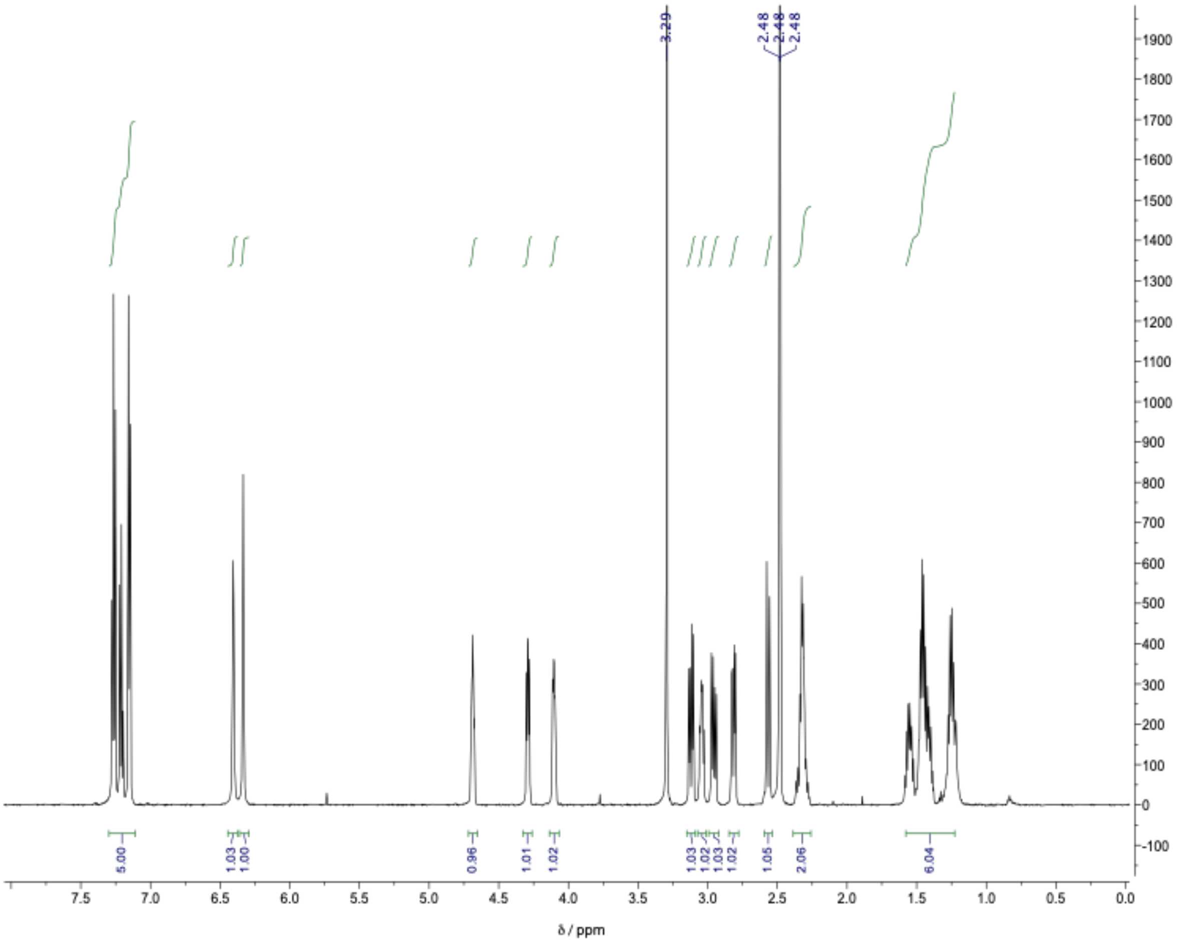

Figure S9, continued.
Biotinyl-leucine oxazolone (BLO)

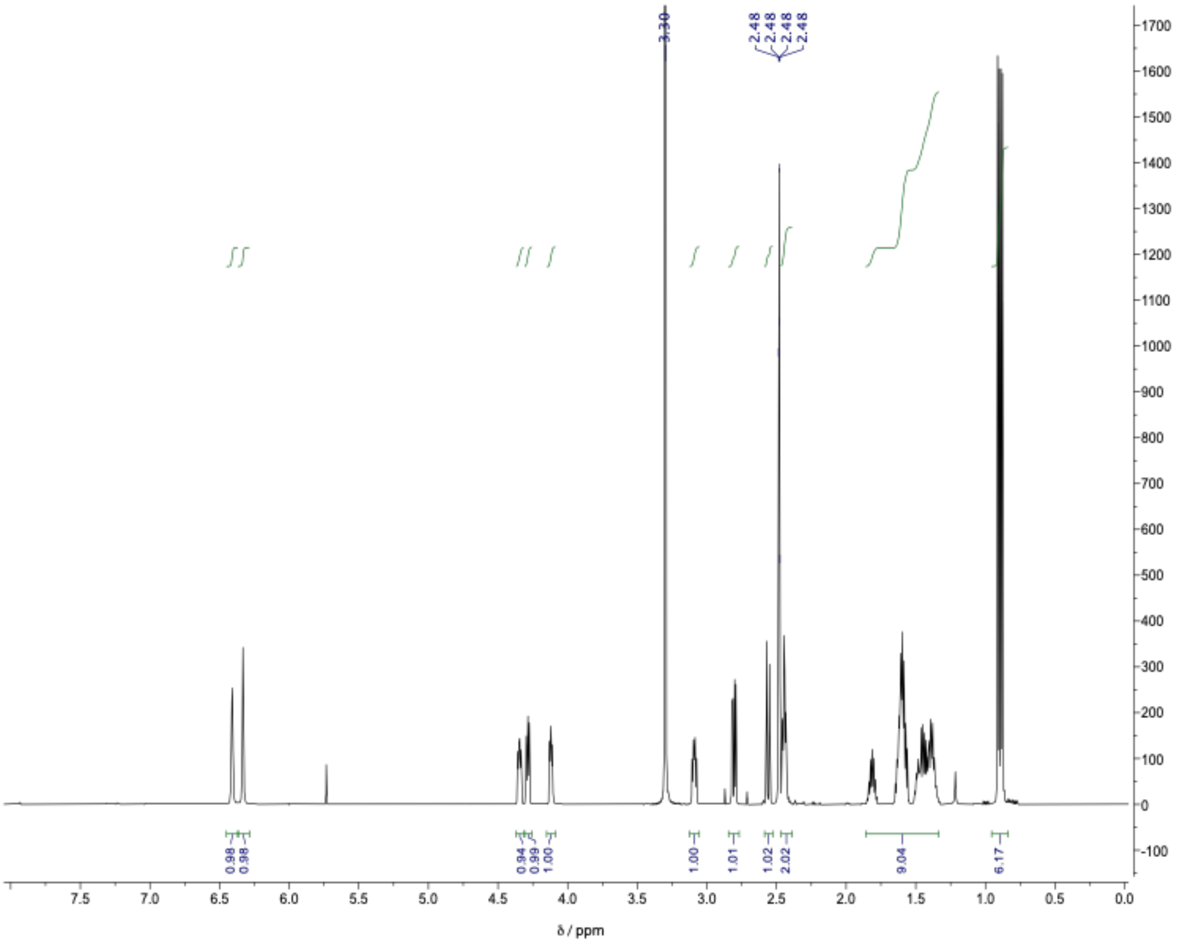

Figure S9, continued.
Biotinyl-isoleucine oxazolone (BIO)

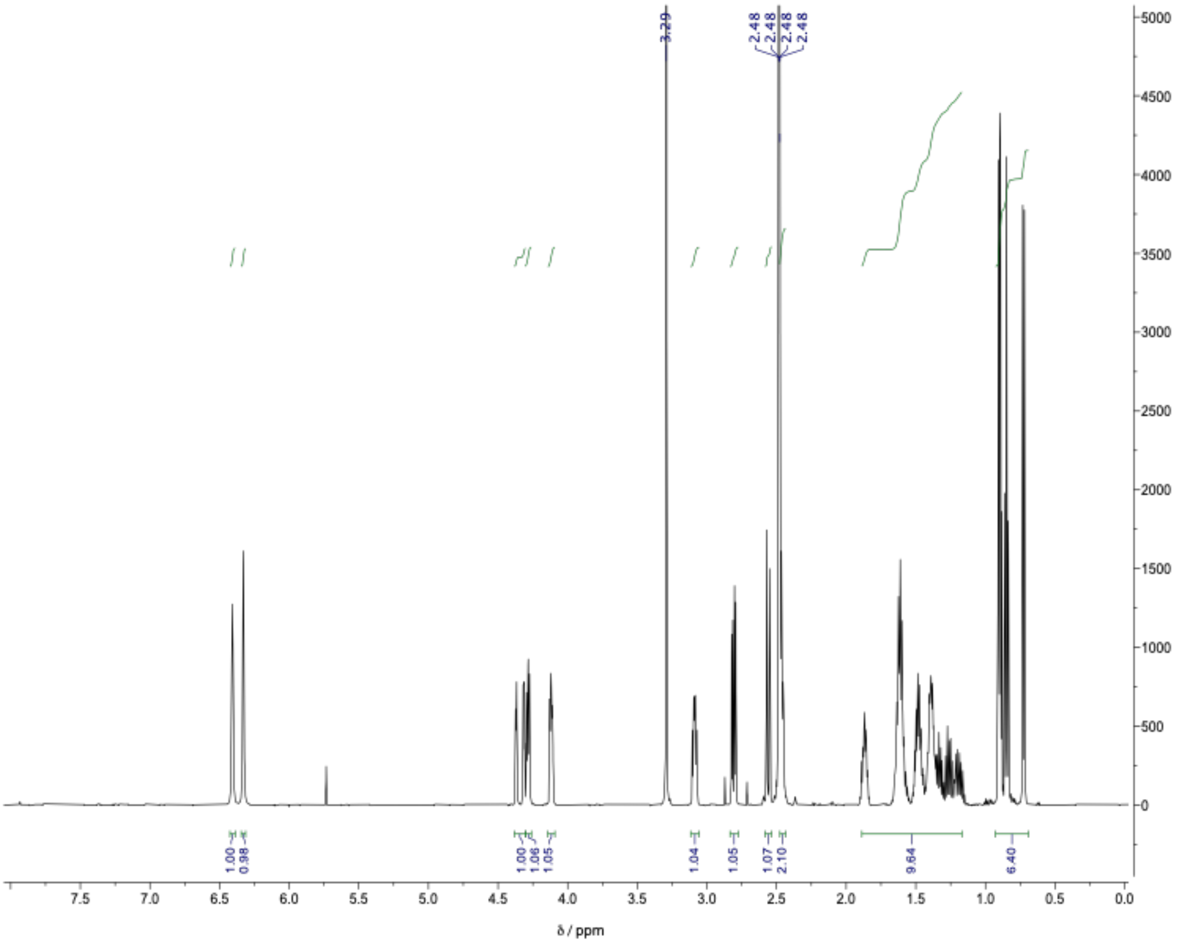

Figure S9, continued.
Biotinyl-valine oxazolone (BVO)

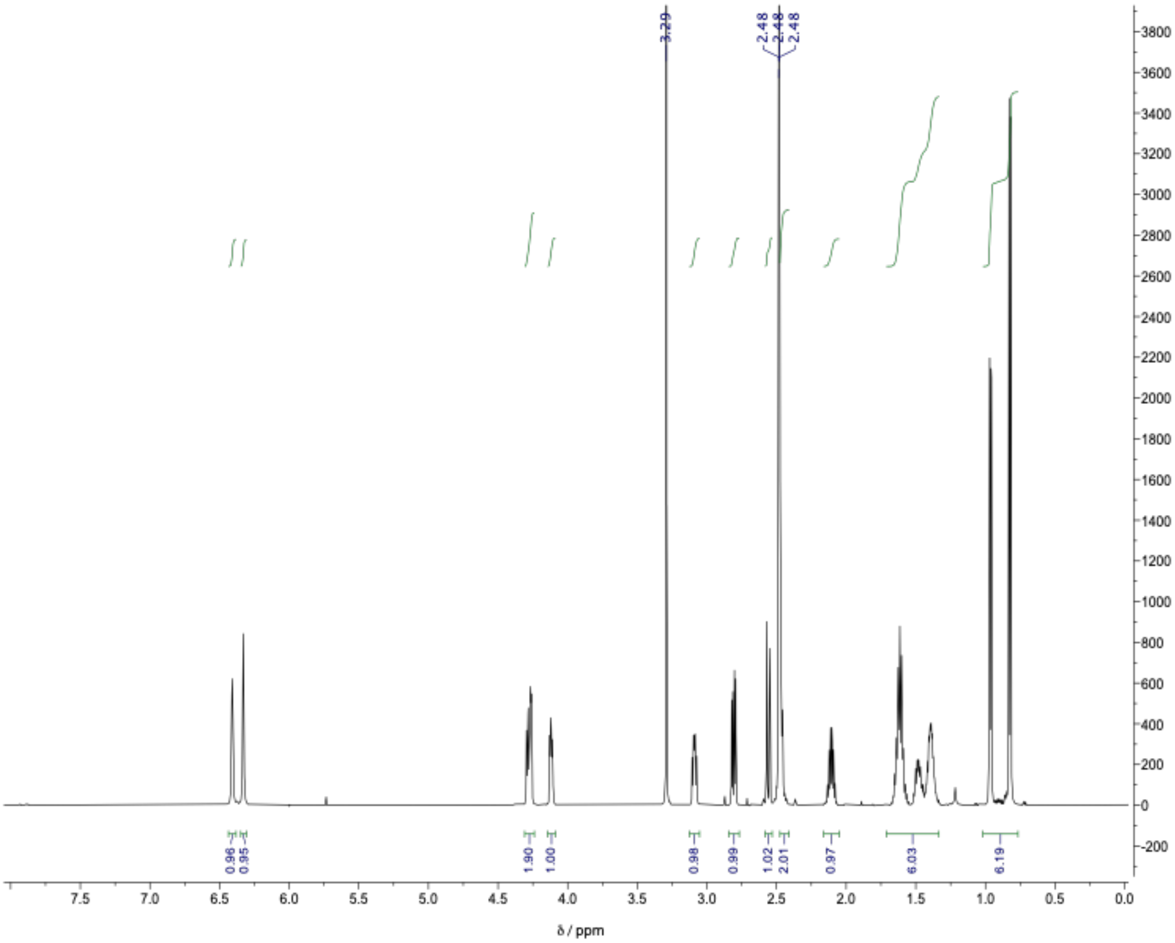

Figure S9, continued.
Biotinyl-methionine oxazolone (BMO)

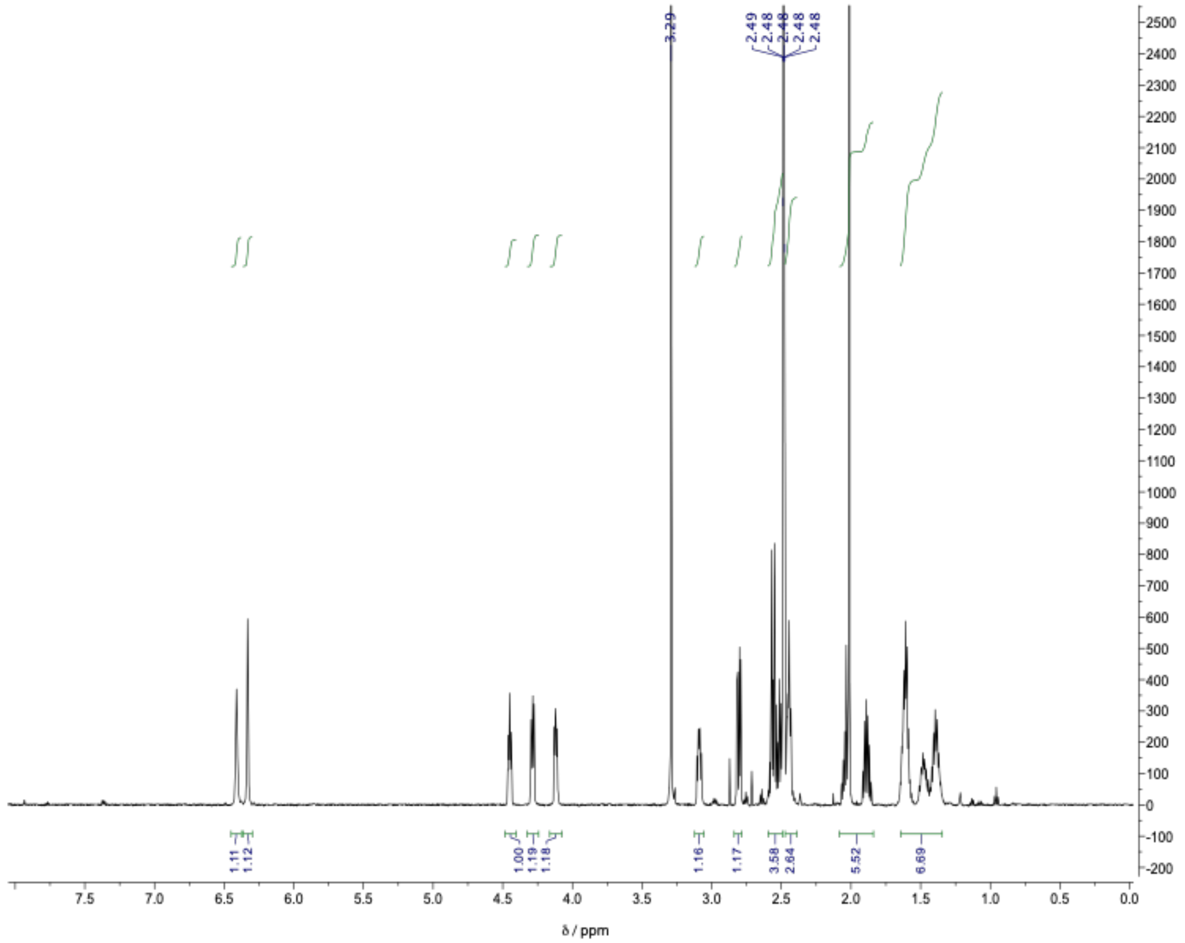

**Table S5.** Biotin quantification of BXO compounds.

| Compound | $[biotin]$ (mM) | σ |
|---|---|---|
| BWO | 9.57 | 0.98 |
| BFO | 10.37 | 1.88 |
| BLO | 14.35 | 0.42 |
| BIO | 13.49 | 1.34 |
| BVO | 14.00 | 2.13 |
| BMO | 12.56 | 1.51 |

**Figure S10**. Representative standard curve for qPCR. Standards are used for calculating [cDNA] from recovered fractions of aminoacylated RNA from *k*-Seq experiments. Shown is $log_{10}$-transformed [ssDNA] (pg/uL) from prepared serial dilutions of Qubit-quantified library ssDNA and the associated quantification cycle ($C_q$) from qPCR. Error bars indicate standard deviation from 15 replicates (five sets of triplicate experiments). The PCR efficiency is estimated to be 97.61%.

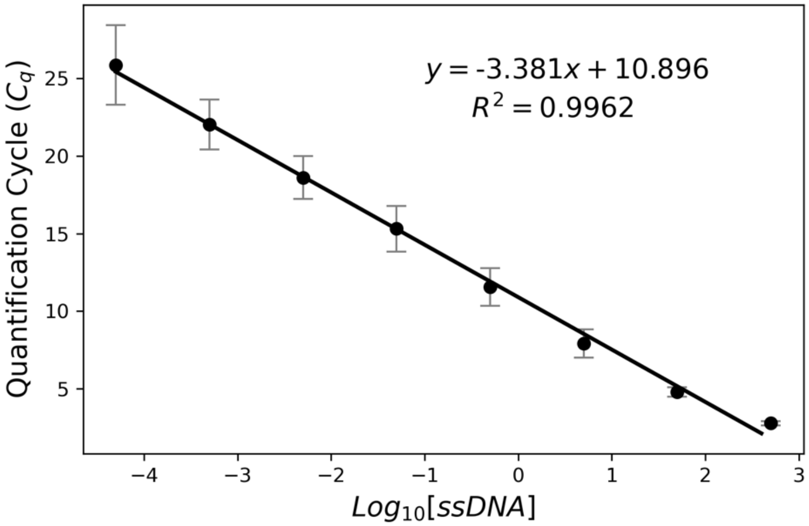
